# PDGFRα/β heterodimer activation negatively affects downstream ERK1/2 signaling and cellular proliferation

**DOI:** 10.1101/2023.12.27.573428

**Authors:** Maria B. Campaña, Madison R. Perkins, Maxwell C. McCabe, Andrew Neumann, Eric D. Larson, Katherine A. Fantauzzo

## Abstract

The platelet-derived growth factor receptor (PDGFR) family of receptor tyrosine kinases allows cells to communicate with one another by binding to growth factors at the plasma membrane and activating intracellular signaling pathways to elicit responses such as migration, proliferation, survival and differentiation. The PDGFR family consists of two receptors, PDGFRα and PDGFRβ, that dimerize to form PDGFRα homodimers, PDGFRα/β heterodimers and PDGFRβ homodimers. Here, we overcame prior technical limitations in visualizing and purifying PDGFRα/β heterodimers by generating a cell line stably expressing C-terminal fusions of PDGFRα and PDGFRβ with bimolecular fluorescence complementation fragments corresponding to the N-terminal and C-terminal regions of the Venus fluorescent protein, respectively. We found that these receptors heterodimerize relatively quickly in response to PDGF-BB ligand treatment, with a peak of receptor autophosphorylation following 5 minutes of ligand stimulation. Moreover, we demonstrated that PDGFRα/β heterodimers are rapidly internalized into early endosomes, particularly signaling endosomes, where they dwell for extended lengths of time. We showed that PDGFRα/β heterodimer activation does not induce downstream phosphorylation of ERK1/2 and significantly inhibits cell proliferation. Further, we characterized the PDGFR dimer-specific interactome and identified MYO1D as a novel protein that preferentially binds PDGFRα/β heterodimers. We demonstrated that knockdown of MYO1D leads to retention of PDGFRα/β heterodimers at the plasma membrane, resulting in increased phosphorylation of ERK1/2 and increased cell proliferation. Collectively, our findings impart valuable insight into the molecular mechanisms by which specificity is introduced downstream of PDGFR activation to differentially propagate signaling and generate distinct cellular responses.

**One-sentence summary:** PDGFRα/β heterodimer binding to MYO1D contributes to rapid internalization of the dimerized receptors, thereby negatively affecting downstream ERK1/2 signaling and cellular proliferation.

## INTRODUCTION

The platelet-derived growth factor receptor (PDGFR) family of receptor tyrosine kinases (RTKs) allows cells to communicate with one another by binding to growth factors at the plasma membrane and activating intracellular signaling pathways to elicit responses such as migration, proliferation, survival and differentiation (*1, 2*). The mammalian PDGF family is composed of four dimeric ligands, PDGF-AA, PDGF-BB, PDGF-CC and PDGF-DD, which variously signal through two receptors, PDGFRα and PDGFRβ. These receptors possess an extracellular ligand-binding domain harboring five immunoglobulin-like loops, a single transmembrane domain and an intracellular domain containing a split, catalytic tyrosine kinase (*3*). PDGFRα and PDGFRβ share the highest amino acid homology in the kinase domains, with significantly reduced homology in the extracellular and interkinase domains (*4*). Upon ligand binding, the PDGFRs dimerize to form PDGFRα homodimers, PDGFRα/β heterodimers or PDGFRβ homodimers (*4–8*). Through an undetermined mechanism, dimerization subsequently promotes tyrosine kinase activity, resulting in the trans-autophosphorylation of intracellular tyrosine residues (*6, 9*). Signaling molecules and adaptor proteins containing Src homology 2 phosphotyrosine recognition motifs subsequently bind to specific phosphorylated residues in the receptors to initiate downstream intracellular signaling cascades (*1, 3*).

Based on *in vivo* mouse knockout phenotypes, the ligands PDGF-AA and PDGF-CC have been shown to solely activate PDGFRα signaling during development, while PDGF-BB activates PDGFRβ signaling (*10–14*). Alternatively, the ligand PDGF-DD is not essential for development nor postnatal life (*15*). While this data is often interpreted as representing the functional roles of PDGFRα homodimers and PDGFRβ homodimers, compelling evidence has emerged indicating that PDGFRα/β heterodimers also form during development. First, analysis of autophosphorylation mutant knockin alleles in mice suggested that PDGFRβ is able to compensate for the loss of PI3K signaling through PDGFRα upon heterodimer formation (*16*). Embryos homozygous for an allele (*Pdgfra^PI3K^*) in which PDGFRα is unable to bind PI3K due to two tyrosine to phenylalanine mutations (*17*) die perinatally and exhibit palatal clefting, among other abnormalities (*8, 16*). However, these embryos do not display the full range of skeletal defects, such as overt facial clefting, observed in *Pdgfra*-null embryos (*11*). In support of PDGFRα/β heterodimer formation, whereas *Pdgfrb^PI3K/PI3K^*mice do not exhibit craniofacial defects (*18*), *Pdgfra^PI3K/PI3K^;Pdgfrb^PI3K/PI3K^* double-homozygous mutant embryos in which PI3K signaling cannot be engaged through PDGFRα/β heterodimers phenocopy *Pdgfra*-null embryos (*16*). More recently, we and others have demonstrated that PDGFRα and PDGFRβ genetically and physically interact in the murine embryonic mesenchyme, primarily in response to PDGF-BB ligand (*8, 19–21*). However, definitive proof of the formation of active PDGFRα/β heterodimers has remained elusive.

Technical limitations had previously prevented an unbiased analysis of PDGFR dimer-specific dynamics, as antibodies cannot differentiate between receptors present as monomers or engaged in homodimers versus heterodimers. Further, ligand identity alone cannot inform dimer-specific conclusions, as some PDGF ligands, especially PDGF-BB, are promiscuous and can result in the formation of multiple dimers (*8*). To overcome these limitations, we recently developed a novel bimolecular fluorescence complementation (BiFC) (*22, 23*) approach that enabled us to visualize and purify activated PDGFRα homodimers and PDGFRβ homodimers (*24*). We cloned plasmids expressing C-terminal fusions of each PDGFR with BiFC fragments corresponding to the N-terminal (V1) or C-terminal (V2) regions of the Venus fluorescent protein (*23*). The individual N- and C-terminal fragments are non-fluorescent when bound to monomeric receptors. However, upon receptor dimerization, the N- and C-terminal fragments colocalize, resulting in a functional Venus protein that can be visualized by fluorescence microscopy or purified biochemically using a GFP-Trap nanobody that has an epitope spanning V1 and V2 (*25*). Using this approach, we demonstrated that PDGFRα receptors homodimerize slowly, have decreased levels of autophosphorylation, are rapidly internalized into early endosomes and are quickly degraded via late endosome trafficking to the lysosome. Alternatively, we found that PDGFRβ receptors homodimerize quickly, have increased levels of autophosphorylation, are slowly internalized and are more likely to be recycled to the plasma membrane via recycling endosomes (*24*). We showed that these dynamics lead to a greater amplitude of ERK1/2 and AKT phosphorylation downstream of PDGFRβ homodimer activation, in addition to increased proliferation and migration (*24*). While these findings provided considerable insight into mechanisms by which specificity is introduced downstream of PDGFR activation, the protein interactions that mediate the internalization and trafficking of the various PDGFR dimers are incompletely understood.

Here, we employed the same BiFC approach to examine the dimerization, activation, internalization and trafficking dynamics of PDGFRα/β heterodimers, as well as their effect on intracellular signaling and cellular behavior, revealing significant differences from what we previously observed for the PDGFR homodimers (*24*). Further, we generated the first PDGFR dimer-specific interactome, identifying proteins that differentially bind the three PDGFR dimers and contribute to receptor internalization following activation at the plasma membrane. Combined, these findings substantially expand our understanding of the mechanisms by which the various PDGFR dimers propagate downstream signaling to generate distinct cellular outputs.

## RESULTS

### Generation and validation of a PDGFRα/β-BiFC stable cell line

To investigate PDGFRα/β-specific dynamics, and compare these to what we previously observed for PDGFRα homodimers and PDGFRβ homodimers (*24*), we stably integrated PDGFRα-V1 and PDGFRβ-V2 sequences (Fig. S1A) into the human-derived HCC15 cell line via lentiviral transduction. We chose this cell line due to its mesenchymal expression profile, lack of *PDGFRA* and *PDGFRB* expression, minimal expression of the various PDGF ligands (with only *PDGFA* being expressed at a low reads per kilobase million (RPKM) value of 2.70), and a spread out morphology that allows for examination of trafficking events (*26, 27*). PDGFRα-V1 and PDGFRβ-V2 sequence integration was confirmed by PCR amplification and Sanger sequencing. Quantitative real-time PCR (qRT-PCR) revealed that the transcripts encoding *PDGFRA* (1.48±0.247% of *B2M* expression; mean±s.e.m.) and *PDGFRB* (1.14±0.365% of *B2M* expression) were expressed at similar levels (Fig. S1B). Lastly, we demonstrated that both PDGFRα and PDGFRβ proteins were expressed in the PDGFRα/β heterodimer cell line and migrated at the expected masses via western blotting (Fig. S1C).

To confirm the Venus complementation event upon exogenous ligand treatment, we starved the cells for 24 h in HITES medium lacking any growth factors, photobleached and stimulated the cells with PDGF-BB ligand for 5 min. We had previously demonstrated that PDGFRα/β heterodimers form most robustly in response to PDGF-BB ligand treatment of both primary mouse embryonic fibroblasts (MEFs) and mouse embryonic palatal mesenchyme (MEPM) cells (*8, 20*). We observed little to no Venus expression in the absence of ligand treatment (Fig. 1A) and robust Venus expression surrounding the nucleus and at the periphery of the cell following ligand stimulation (Fig. 1B). Quantification of fluorescence intensity demonstrated a significant increase in Venus intensity between no ligand treatment (6.61±0.264 arbitrary units (A.U.); mean±s.e.m.) and 5 min of PDGF-BB treatment (34.9±4.51 A.U.) (Fig. 1C). We next examined colocalization of the Venus signal with signals from anti-PDGFRα and anti-PDGFRβ antibodies. In each case, Venus signal colocalized with a subset of PDGFR expression (Fig. 1D to F). The extent of colocalization of the Venus signal with signals from anti-PDGFRα (0.346±0.0359 Pearson’s correlation coefficient (PCC); mean±s.e.m.) and anti-PDGFRβ (0.359±0.0678 PCC) antibodies was comparable in the PDGFRα/β heterodimer cell line (Fig. 1D). The observed PCC values less than 1 indicated that the presence of the BiFC fragments alone was not driving heterodimerization (Fig. 1D). The regions of receptor expression that were not Venus-positive (Fig. 1E and F) likely represent monomeric receptors and/or potential PDGFRαV1/αV1 and especially PDGFRβV2/βV2 homodimers, which would not be expected to fluoresce.

**Fig. 1.**
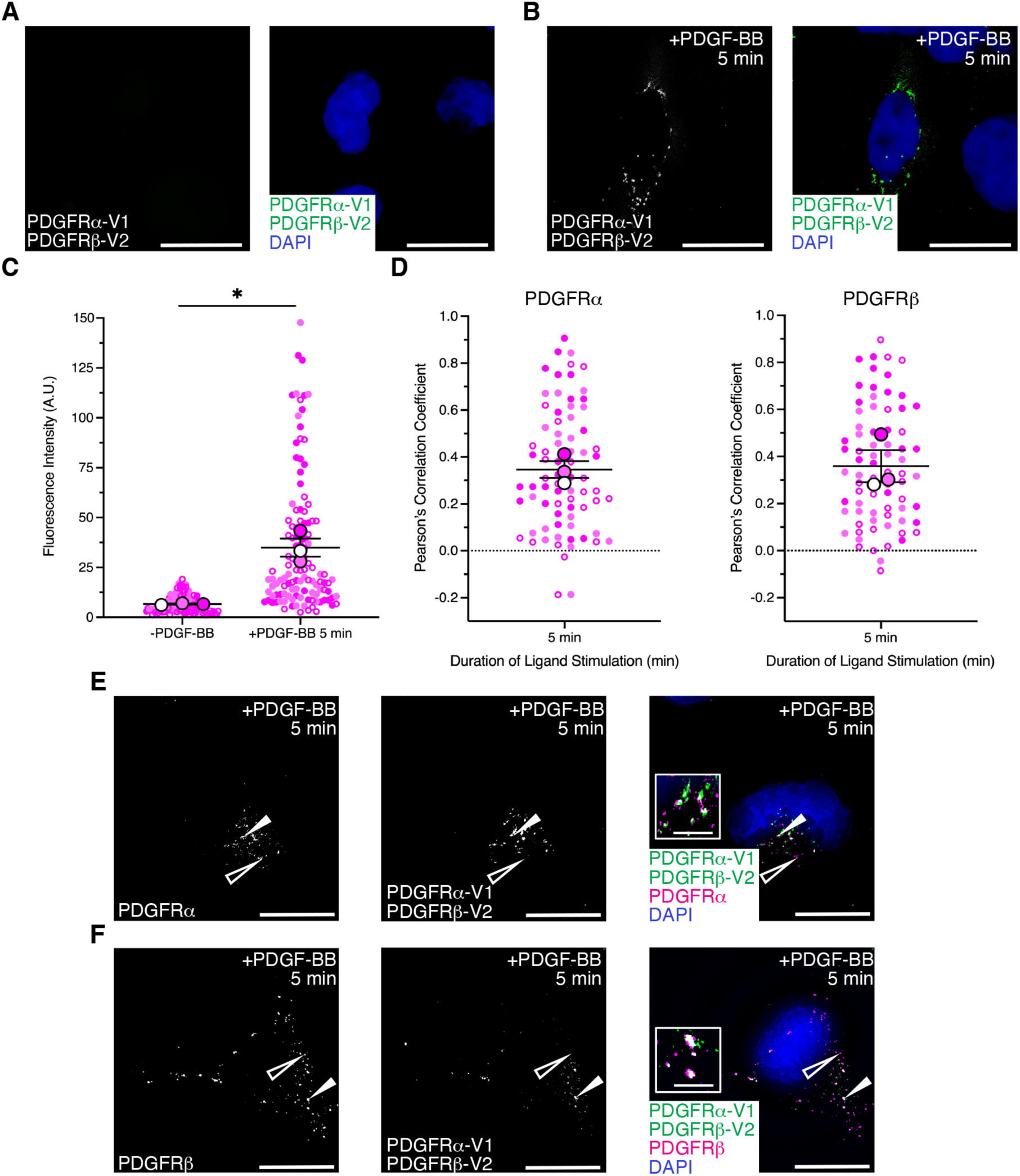
Validation of a PDGFRα/β-BiFC stable cell line. (**A,B**) Venus expression (white or green) as assessed by fluorescence analysis of HCC15 cells transduced with PDGFRα-V1 and PDGFRβ-V2 in the absence (A) or presence (B) of PDGF-BB ligand for 5 min. Nuclei were stained with DAPI (blue; A,B). Scale bars: 20 µm. (**C**) Scatter dot plot depicting fluorescence intensity for the PDGFRα/β heterodimer cell line in the absence or presence of PDGF-BB ligand for 5 min. Data are mean±s.e.m. \**P*<0.05 (two-tailed, paired *t*-test). Colored circles correspond to independent experiments. Summary statistics from biological replicates consisting of independent experiments (large circles) are superimposed on top of data from all cells; *n*≥38 technical replicates across each of three biological replicates. (**D**) Scatter dot plots depicting Pearson’s correlation coefficient of the PDGFRα/β heterodimer cell line with an anti-PDGFRα antibody (left) and an anti-PDGFRβ antibody (right) following PDGF-BB ligand stimulation for 5 min. Data are mean±s.e.m. Colored circles correspond to independent experiments. Summary statistics from biological replicates consisting of independent experiments (large circles) are superimposed on top of data from all cells; *n*=25 technical replicates across each of three biological replicates. (**E,F**) PDGFRα (E) and PDGFRβ (F) antibody signal (white or magenta; E,F) and/or Venus expression (white or green; E,F) as assessed by (immuno)fluorescence analysis of the PDGFRα/β heterodimer cell line. Insets in E and F are regions where white arrows are pointing. Nuclei were stained with DAPI (blue; E,F). White arrows denote colocalization; white outlined arrows denote lack of colocalization. Scale bars: 20 µm (main images), 3 µm (insets).

### PDGFRα/β heterodimers dimerize relatively quickly and have peak levels of autophosphorylation following 5 minutes of ligand treatment

To confirm the Venus complementation event upon exogenous ligand treatment biochemically, we stimulated the cells with PDGF-BB ligand for 2 and 5 min. Lysates were immunoprecipitated with the GFP-Trap nanobody and subjected to western blotting for PDGFRα and PDGFRβ, normalizing the immunoprecipitation signal to the whole-cell lysate signal for each receptor. We found that the receptors heterodimerize relatively quickly, with peak dimerization following 5 min of ligand treatment for both PDGFRα (2.17±1.27 relative induction (R.I.); mean±s.e.m.) and PDGFRβ (1.95±0.173 R.I.) (Fig. 2A and B). As would be expected for heterodimers, the relative induction values for the two receptors were essentially equal throughout the time course (Fig. 2B). We next examined activation of the receptors by assessing receptor autophosphorylation. Cells were stimulated with PDGF-BB ligand in a time course from 2 min to 4 h. Whole-cell lysates were immunoprecipitated with the GFP-Trap nanobody and subjected to western blotting with an anti-phospho-PDGFR antibody, which recognizes a phosphorylated tyrosine residue within the tyrosine kinase domains of both PDGFRα and PDGFRβ. Consistent with previous findings demonstrating that both ligand binding and dimerization are required for receptor activation (*9, 24, 28*), we observed minimal phospho-PDGFR signal in the absence of ligand stimulation (Fig. 2C). However, the receptors were significantly autophosphorylated in response to ligand treatment, with peak activation following 5 min of PDGF-BB stimulation (6.61±1.75 R.I.) (Fig. 2C and D). As we previously observed, the anti-PDGFR blots contained two bands representing the non-glycosylated and glycosylated forms of the receptors (*7, 24*), while the anti-phospho-PDGFR blots possessed a single band representing the glycosylated, phosphorylated versions of the receptors upon ligand treatment (*24*).

**Fig. 2.**
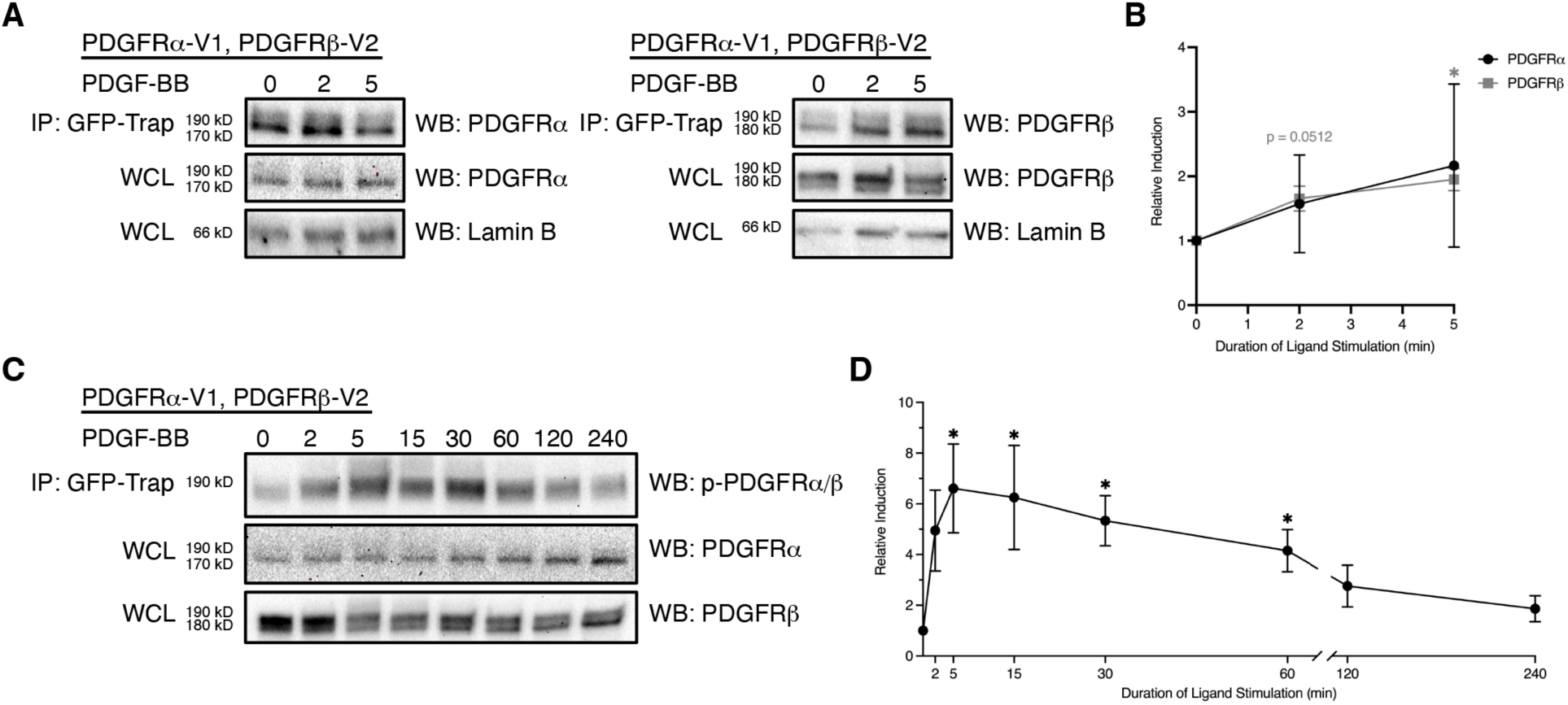
PDGFRα/β heterodimers dimerize relatively quickly and have peak levels of autophosphorylation following 5 minutes of ligand treatment. (**A**) Immunoprecipitation (IP) of dimerized PDGFRα/β receptors with GFP-Trap nanobody from cells that were unstimulated or treated with PDGF-BB ligand for 2–5 min followed by western blotting (WB) with an anti-PDGFRα (left) or an anti-PDGFRβ antibody (right). WCL, whole-cell lysates. (**B**) Line graph depicting quantification of band intensities from *n*=3 biological replicates as in A. Data are mean±s.e.m. \**P*<0.05 (two-tailed, ratio paired *t*-test). (**C**) Immunoprecipitation of dimerized PDGFRα/β receptors with GFP-Trap nanobody from cells that were unstimulated or treated with PDGF-BB ligand for 2 min to 4 h followed by western blotting with an anti-phospho (p)-PDGFR antibody. (**D**) Line graph depicting quantification of band intensities from *n*=3 biological replicates as in C. Data are mean±s.e.m. \**P*<0.05; (two-tailed, ratio paired *t*-test).

### PDGFRα/β heterodimers are rapidly internalized into signaling endosomes

Following dimerization and activation at the cell membrane in response to ligand binding, PDGFRs are internalized and trafficked through the early endosome before being trafficked to the lysosome via late endosomes for degradation or recycled to the membrane via recycling endosomes for continued signaling (*24, 29–33*). To investigate the trafficking dynamics of PDGFRα/β heterodimers in response to ligand treatment, we examined colocalization of the Venus signal in our PDGFRαV1/βV2 cell line with markers of various subcellular compartments. We found that PDGFRα/β heterodimers were rapidly internalized, as indicated by a significant decrease in colocalization with the plasma membrane marker Na^+^/K^+^-ATPase (*34*) from the 1 min (0.217±0.0134 PCC) to 5 min (0.068±0.00956 PCC) ligand treatment timepoints (Fig. 3A to C). Further, analysis of Venus colocalization with the early endosome marker Rab5 (all isoforms) (*35*) demonstrated that PDGFRα/β heterodimers were internalized into early endosomes, where they dwelled at relatively stable levels from 2 min (0.446±0.0401 PCC) to 30 min (0.404±0.0503 PCC) following ligand treatment (Fig. 3D to F). PDGFRα/β heterodimers were primarily found in signaling endosomes during this time, as colocalization values were higher for the signaling endosome marker APPL1 (*36*) at 2 min (0.193±0.0468 PCC) and 30 min (0.245±0.0476 PCC) of ligand treatment (Fig. 3G to I) than the non-signaling endosome marker EEA1 (*37*) at these same timepoints (0.0932±0.0116 PCC and 0.0344±0.0136 PCC, respectively) (Fig. S2A to C). Following trafficking to the early endosomes, the PDGFRα/β heterodimers were subsequently trafficked to RAB7-positive late endosomes (all isoforms) (*35*) (Fig. S2D and E), RAB4-positive rapid recycling endosomes (all isoforms) (*35*) (Fig. S2F and G) and RAB11-positive slow recycling endosomes (all isoforms) (*35*) (Fig. S2H and I). Colocalization of Venus signal with Na^+^/K^+^-ATPase at 60 min (0.199±0.0132 PCC) and 90 min (0.178±0.0115 PCC) of ligand treatment (Fig. S2J and K) was comparable to that following 1 min of ligand treatment described above (Fig. 3A and B), suggesting that a subset of PDGFRα/β heterodimers are recycled following ligand stimulation. Collectively, these data indicate that PDGFRα/β heterodimers are rapidly internalized into signaling endosomes, where they dwell for extended lengths of time, before being trafficked for degradation and recycling.

**Fig. 3.**
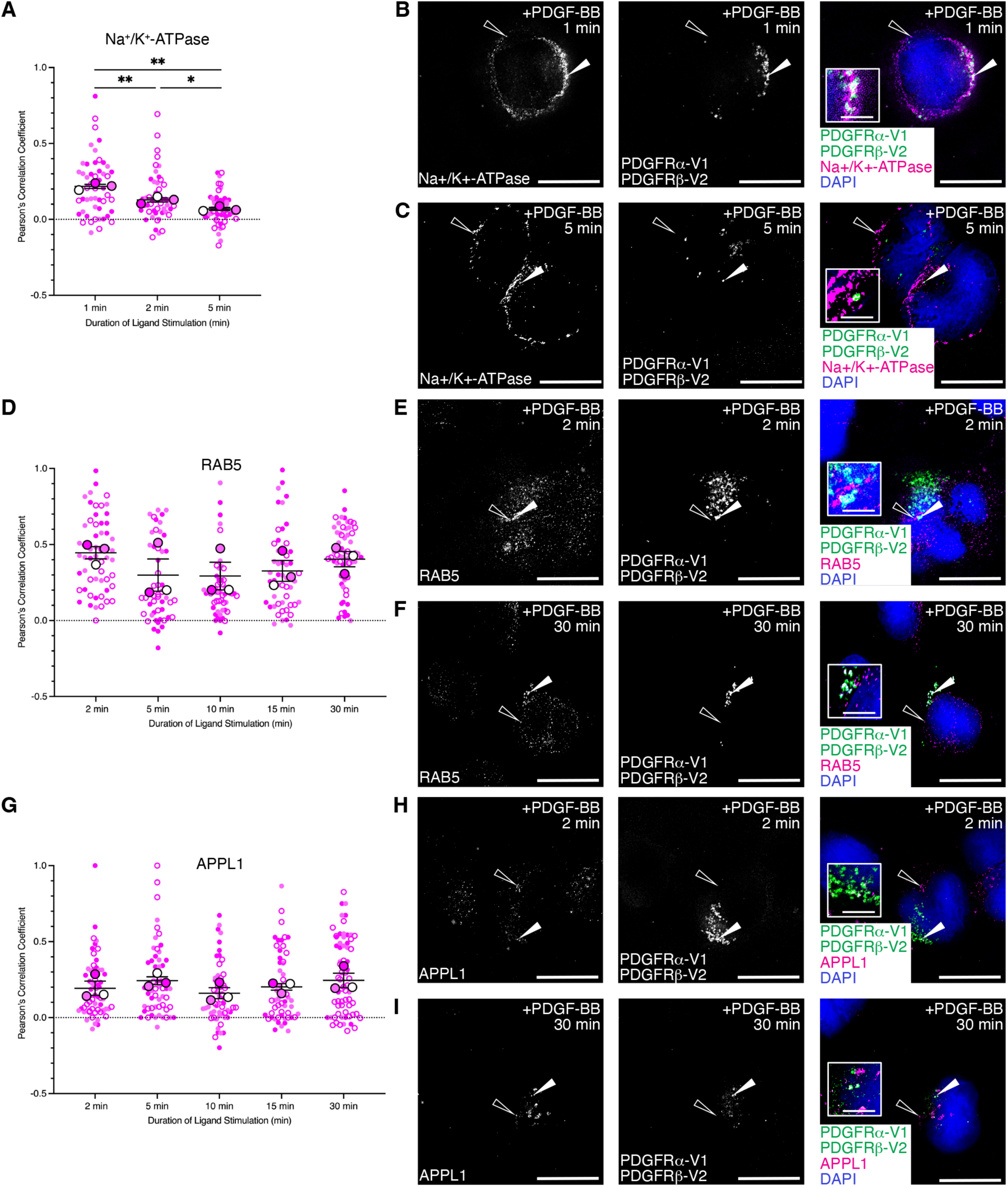
PDGFRα/β heterodimers are rapidly internalized into signaling endosomes. (**A,D,G**) Scatter dot plots depicting Pearson’s correlation coefficient of the PDGFRα/β heterodimer cell line Venus signal with an anti-Na^+^/K^+^-ATPase antibody (A), an anti-RAB5 antibody (D) or an anti-APPL1 antibody (G) signal following PDGF-BB ligand stimulation from 1–5 min (A) or 2-30 min (D,G). Data are mean±s.e.m. \**P*<0.05; \*\**P*<0.01 (two-tailed, unpaired *t*-test with Welch’s correction). Colored circles correspond to independent experiments. Summary statistics from biological replicates consisting of independent experiments (large circles) are superimposed on top of data from all cells; *n*≥11 technical replicates across each of three biological replicates. (**B,C,E,F,H,I**) Na^+^/K^+^-ATPase antibody signal (white or magenta; B,C), RAB5 antibody signal (white or magenta; E,F) or APPL1 antibody signal (white or magenta; H,I) and/or Venus expression (white or green; B,C,E,F,H,I) as assessed by (immuno)fluorescence analysis of the PDGFRα/β heterodimer cell line. Insets in B, C, E, F, H and I are regions where white arrows are pointing. Nuclei were stained with DAPI (blue; B,C,E,F,H,I). White arrows denote colocalization; white outlined arrows denote lack of colocalization. Scale bars: 20 µm (main images), 3 µm (insets).

### PDGFRα/β heterodimer activation does not induce downstream phosphorylation of ERK1/2 and inhibits cell proliferation

To examine the effects of the above PDGFRα/β heterodimer-specific trafficking dynamics on downstream intracellular signaling, we performed a time course of ligand stimulation from 2 min to 4 h followed by western blotting of whole-cell lysates for phosphorylation of two effector molecules downstream of PDGFR activation, ERK1/2 and AKT (all isoforms) (*8, 24, 38*). PDGFRα/β heterodimer activation did not induce downstream phosphorylation of ERK1/2, as relative induction levels did not increase beyond baseline levels at 0 min of ligand stimulation (1.00±0.00 R.I.) until 4 h of PDGF-BB treatment (1.08±0.506 R.I.) (Fig. 4A and B). Alternatively, PDGFRα/β heterodimer activation led to a very transient phospho-AKT response, with a peak at 5 min of ligand stimulation (2.74±0.202 R.I.) and a return to near baseline levels by 30 min of PDGF-BB treatment (1.27±0.366 R.I.) that remained relatively stable throughout the rest of the time course (1.25±0.194 R.I. at 4 h) (Fig. 4C and D).

**Fig. 4.**
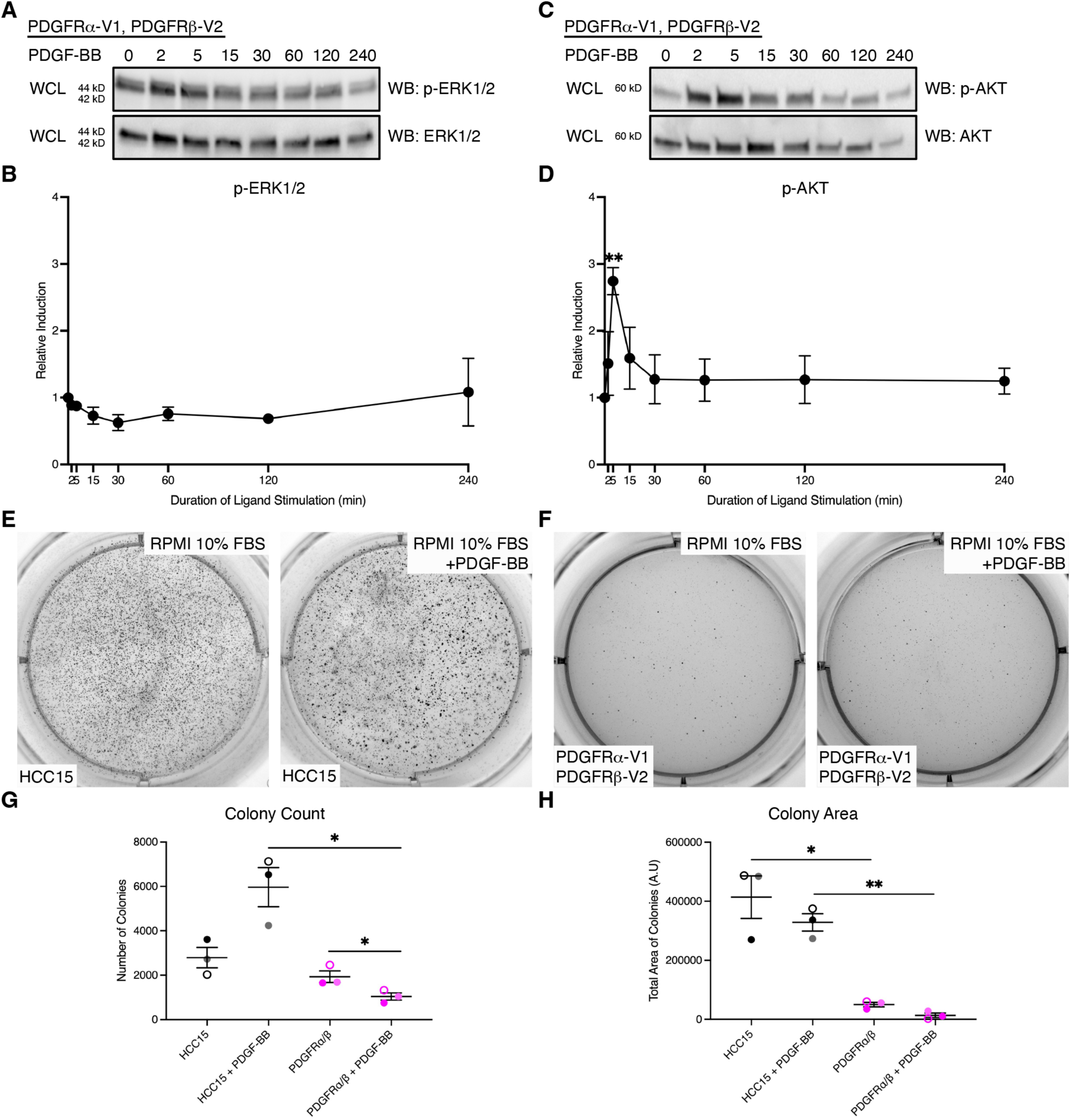
PDGFRα/β heterodimer activation does not induce downstream phosphorylation of ERK1/2 and inhibits cell proliferation. (**A,C**) Western blot (WB) analysis of whole-cell lysates (WCL) from the PDGFRα/β heterodimer cell line following a time course of PDGF ligand stimulation from 2 min to 4 h with anti-phospho (p)-ERK1/2 (A) or anti-phospho (p)-AKT (C) antibodies. (**B,D**) Line graphs depicting quantification of band intensities from *n*=3 biological replicates as in A and C. Data are mean±s.e.m. \*\**P*<0.01 (two-tailed, ratio paired *t*-test). (**E,F**) Colony growth in soft agar anchorage-independent growth assays for the HCC15 cell line (E) or the PDGFRα/β heterodimer cell line (F) after 10 days in RPMI growth medium in the absence or presence of PDGF-BB ligand in six-well plate wells. (**G,H**) Scatter dot plots depicting quantification of colony count (G) or colony area (H) from *n*=3 biological replicates as in E and F. Data are mean±s.e.m. \**P*<0.05; \*\**P*<0.01 (two-tailed, paired *t-*test within each cell line and a two-tailed, unpaired *t*-test with Welch’s correction between each cell line). Colored symbols correspond to independent experiments.

Proliferation is increased downstream of both PDGFRα and PDGFRβ signaling in settings where both receptors are active (*21, 39, 40*). To determine how the above trafficking and signaling dynamics affect cellular activity downstream of PDGFRα/β heterodimer activation, we assessed cell proliferation via a soft agar anchorage-independent growth assay in RPMI growth medium with 10% fetal bovine serum (FBS) over the course of 10 days. As expected, HCC15 cells, which do not express PDGFRs, did not have significantly increased colony count nor colony area upon PDGF-BB ligand stimulation (Fig. 4E,G and H). In the absence and presence of PDGF-BB ligand, parental HCC15 cells grew visible colonies by the end of the assay (Fig. 4E), with 2.97×10^3^±4.59×10^2^ colonies (mean±s.e.m.) and 5.97×10^3^±8.80×10^2^ colonies (Fig. 4G), and 4.14×10^5^±7.21×10^4^ A.U. and 3.29×10^5^±2.96×10^4^ A.U. total colony area (Fig. 4H), respectively. PDGFRαV1/βV2 cells, however, proliferated less than parental HCC15 cells, and significantly so in the presence of PDGF-BB ligand (Fig. 4F), with 1.04×10^3^±1.64×10^2^ colonies (Fig. 4G) and 1.38×10^4^±7.52×10^3^ A.U. total colony area (Fig. 4H).

Given that PDGFRβ homodimers and PDGFRα/β heterodimers bind the same ligand, PDGF-BB, together, these findings indicated that PDGFRα/β heterodimers may serve as sinks that bind PDGF-BB and prevent the ligand from activating PDGFRβ homodimers. To test this hypothesis, we first demonstrated that PDGF-BB ligand could activate PDGFRβ homodimers in our PDGFRαV1/βV2 cell line. We stimulated PDGFRαV1/βV2 cells with PDGF-BB ligand, resulting in the potential formation of both PDGFRα/β heterodimers and PDGFRβ homodimers. Following immunoprecipitation with the GFP-Trap nanobody, which should only pull down PDGFRα/β heterodimers, the supernatant was immunoprecipitated with an anti-PDGFRβ antibody, which should isolate PDGFRβ monomers and PDGFRβ homodimers. Each sample was then subjected to western blotting with the anti-phospho-PDGFR antibody to identify activated dimers (Fig. S3A). This experiment revealed that both active PDGFRα/β heterodimers and PDGFRβ homodimers formed (Fig. S3B). However, there was only a 4.41±1.56 R.I. for PDGFRβ homodimers following 5 min of ligand stimulation in the PDGFRαV1/βV2 cell line (Fig. S3C), compared to a 26.8±2.04 R.I. that we previously observed at this same timepoint in the PDGFRβV1/βV2 cell line (*24*), indicating that PDGFRα/β heterodimers may indeed serve as ligand sinks.

### MYO1D preferentially binds PDGFRα/β heterodimers

We next combined BiFC with stable isotope labeling by amino acids in cell culture (SILAC) and quantitative mass spectrometry to identify proteins that differentially interact with the various PDGFR dimers at the time of internalization. We used a control cell line with myristoylated, membrane-targeted Venus, as well as our PDGFRαV1/αV2 (*24*), PDGFRαV1/βV2 and PDGFRβV1/βV2 (*24*) cell lines representing each of the three PDGFR dimers. Cells were either grown in the presence of medium with light isotopes and left unstimulated, or grown in the presence of medium with heavy isotopes and stimulated with the relevant PDGF ligand for 5 minutes. Following cell lysis and extraction, we mixed light and heavy samples 1:1 for each cell line, immunoprecipitated the receptor dimers and interacting proteins with the GFP-Trap nanobody, and generated tryptic peptides for mass spectrometry analysis (Fig. 5A). Following total signal normalization and removal of common contaminants (*41*) that were not differentially detected across samples, we filtered the proteins based on three criteria to identify those with increased binding to one or more PDGFR dimer in response to PDGF ligand treatment: 1) the protein had to be present in both mass spectrometry biological replicates for a given cell line, 2) the heavy/light ratio for the experimental sample had to be greater than that for the control myristoylated Venus sample, and 3) the experimental sample heavy/light ratio had to be greater than 1. These parameters generated a list of 350 total proteins, with 177 proteins binding PDGFRα homodimers, and 199 and 217 proteins binding PDGFRα/β heterodimers and PDGFRβ homodimers, respectively. Of these, 71 proteins were shared amongst all three PDGFR dimers, 51 were unique to PDGFRα homodimers, 67 were unique to PDGFRα/β heterodimers, and 60 were unique to PDGFRβ homodimers (Fig. 5B).

**Fig. 5.**
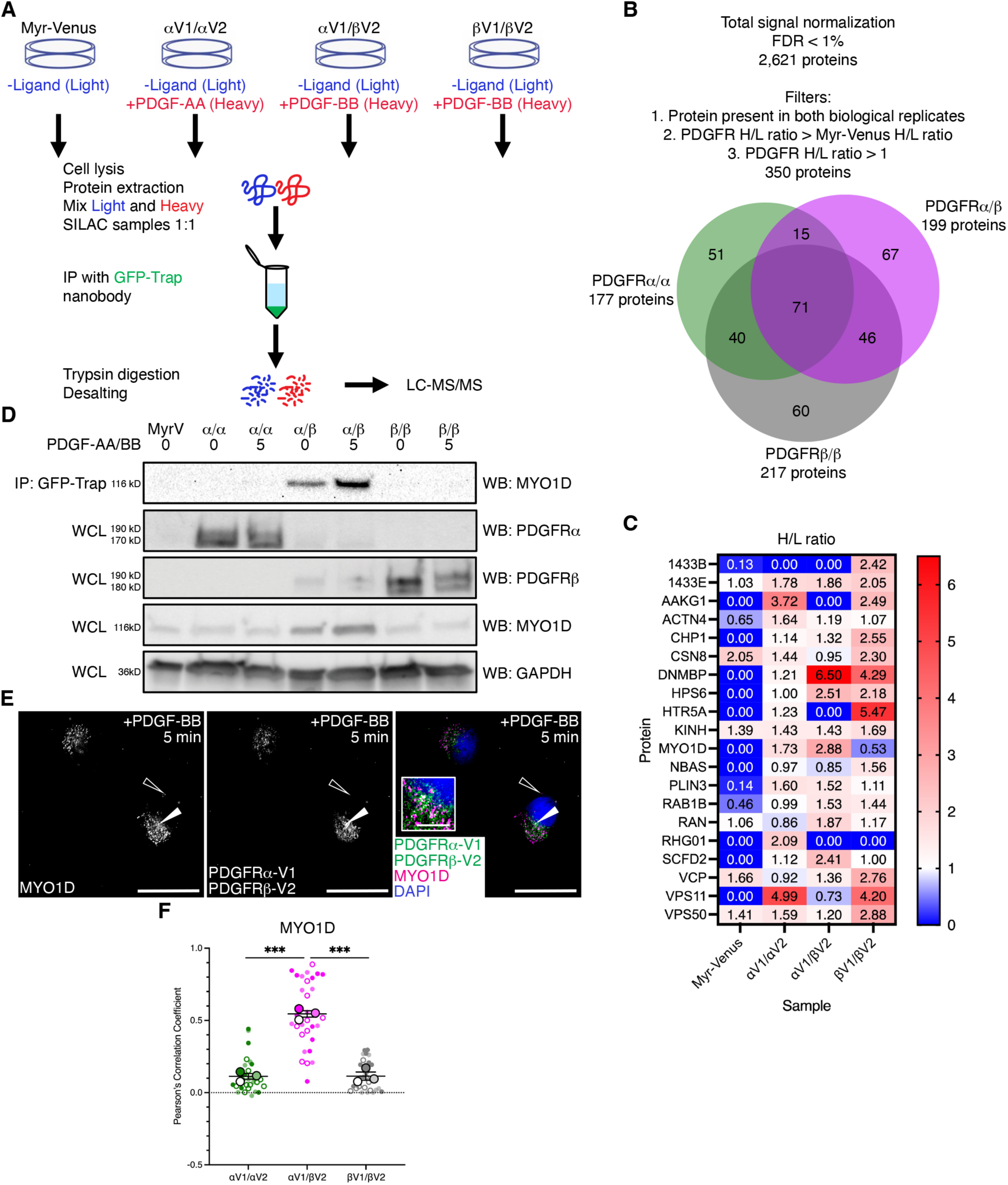
MYO1D preferentially binds PDGFRα/β heterodimers. (**A**) Schematic depicting experimental workflow for identification of PDGFR dimer-specific interacting proteins. (**B**) Filters applied to the mass spectrometry data set and Venn diagram displaying shared protein identifications between samples within the filtered data. (**C**) Heat map of heavy (H)/light (L) SILAC ratios for identified proteins involved in trafficking. (**D**) Immunoprecipitation (IP) of myristoylated Venus and dimerized PDGFR receptors with GFP-Trap nanobody from cells that were unstimulated or treated with PDGF ligand for 5 min followed by western blotting (WB) with an anti-MYO1D antibody. WCL, whole-cell lysates. (**E**) MYO1D antibody signal (white or magenta) and/or Venus expression (white or green) as assessed by (immuno)fluorescence analysis of the PDGFRα/β heterodimer cell line following PDGF-BB ligand stimulation for 5 min. Inset in E is a region where white arrows are pointing. Nuclei were stained with DAPI (blue). White arrows denote colocalization; white outlined arrows denote lack of colocalization. Scale bars: 20 µm (main images), 3 µm (insets). (**F**) Scatter dot plot depicting Pearson’s correlation coefficient of the PDGFRα homodimer, PDGFRα/β heterodimer or PDGFRβ homodimer cell line Venus signal with an anti-MYO1D antibody signal following PDGF ligand stimulation for 5 min as in E. Data are mean±s.e.m. \*\*\**P*<0.001 (two-tailed, unpaired *t*-test with Welch’s correction). Colored circles correspond to independent experiments. Summary statistics from biological replicates consisting of independent experiments (large circles) are superimposed on top of data from all cells; *n*=10 technical replicates across each of three biological replicates.

Known PDGFR-binding proteins were detected in our mass spectrometry screen. For example, the catalytic subunit of PI3K, p110β, was bound to both PDGFRα/β heterodimers and PDGFRβ homodimers, however only in a single biological replicate for each dimer. Gene ontology (GO) analysis with the Enrichr tool (*42*) demonstrated that the most enriched biological process annotations for the 71 proteins common to all three dimers related to mRNA processing and mRNA splicing, as well as anterograde dendritic transport (Fig. S4). The proteins unique to PDGFRα homodimers were additionally enriched for terms related to cytoskeleton organization, regulation of DNA recombination, mitosis, regulation of protein localization to the nucleus and regulation of p53 signal transduction (Fig. S4). The proteins unique to PDGFRα/β heterodimers were additionally enriched for nucleosome assembly and organization, regulation of autophagy, regulation of immunoglobulin production and regulation of gene expression (Fig. S4). Finally, the proteins unique to PDGFRβ homodimers were additionally enriched for retrograde vesicle-mediated transport, regulation of ubiquitin-protein transferase activity, nucleosome disassembly and regulation of microtubule depolymerization and nucleation (Fig. S4), possibly pointing to unique functions for the individual dimers.

Given our demonstration that the various PDGFR dimers have different trafficking dynamics (*24*), we next performed GO analysis of our filtered proteins to identify those involved in trafficking that differentially interact with the various PDGFR dimers (Fig. 5C). While some of these 20 proteins had been implicated in RTK binding, none had previously been reported to interact with the PDGFRs. We chose to focus our initial studies on unconventional myosin-Id (MYO1D), as this protein was not detected in the control myristoylated Venus sample and had differential interaction with the various PDGFR dimers upon ligand stimulation, with the greatest binding to PDGFRα/β heterodimers (Fig. 5C). Further, MYO1D had previously been shown to bind and affect the subcellular localization of the RTKs EGFR, ERBB2 and ERBB4 (*43*). We confirmed that MYO1D preferentially binds PDGFRα/β heterodimers via immunoprecipitation of whole-cell lysates from each cell line with the GFP-Trap nanobody followed by western blotting with an anti-MYO1D antibody (Fig. 5D). Further, these results confirmed our mass spectrometry results that MYO1D exhibits increased binding to PDGFRα/β heterodimers upon 5 min of PDGF-BB ligand treatment compared to binding levels in unstimulated cells (Fig. 5D). Of note, we observed increased MYO1D expression in whole-cell lysates from the PDGFRαV1/βV2 cell line compared to both the PDGFRαV1/αV2 homodimer and PDGFRβV1/βV2 homodimer cell lines (Fig. 5D). Quantitative RT-PCR indicated that this upregulation is observed on the transcript level as well (Fig. S5). This upregulation could stem from a mutation in the clone chosen for the PDGFRαV1/βV2 cell line, disruption of *MYO1D* regulatory elements upon insertion of the PDGFR-BiFC sequences, and/or increased *MYO1D* transcriptional activity or transcript stability upon PDGFRα/β heterodimer activation. In favor of the latter hypotheses, *MYO1D* expression increased with passage number for the PDGFRαV1/βV2 cell line (Fig. S5). Finally, we examined colocalization of MYO1D with the various PDGFR dimers following 5 min of PDGF ligand stimulation via immunofluorescence analysis, demonstrating significantly increased colocalization with PDGFRα/β heterodimers (0.546±0.0424 PCC) (Fig. 5E) compared to either PDGFRα homodimers (0.113±0.0216 PCC) or PDGFRβ homodimers (0.115±0.0183 PCC) (Fig. 5F).

### Knockdown of MYO1D leads to retention of PDGFRα/β heterodimers at the plasma membrane, increased downstream phosphorylation of ERK1/2 and increased cell proliferation

To examine the effects of MYO1D knockdown on PDGFRα/β heterodimer trafficking, we transfected PDGFRαV1/βV2 cells with a Silencer Select negative control siRNA or a combination of two Silencer Select siRNAs targeting *MYO1D* for 48 h. We confirmed efficient knockdown of MYO1D with the siRNAs targeting *MYO1D* via western blotting of whole-cell lysates at this timepoint (Fig. S6). We first assessed colocalization of the Venus signal with Na^+^/K^+^-ATPase, demonstrating a significant increase in colocalization upon treatment with siRNAs targeting *MYO1D* (0.164±0.0227 PCC) compared to the negative control (0.0728±0.00644 PCC) following 5 min of ligand stimulation (Fig. 6A to C). Accordingly, we observed a significant decrease in colocalization with RAB5 upon treatment with siRNAs targeting *MYO1D* (0.123±0.0247 PCC) compared to the negative control (0.315±0.0107 PCC) at 2 min of ligand stimulation (Fig. 6D to F). We did not detect significant differences in colocalization with RAB7 between the two treatments at 1 h of ligand stimulation (Fig. 6G to I), indicating that knockdown of MYO1D primarily effects early internalization and trafficking dynamics of the PDGFRα/β heterodimers.

**Fig. 6.**
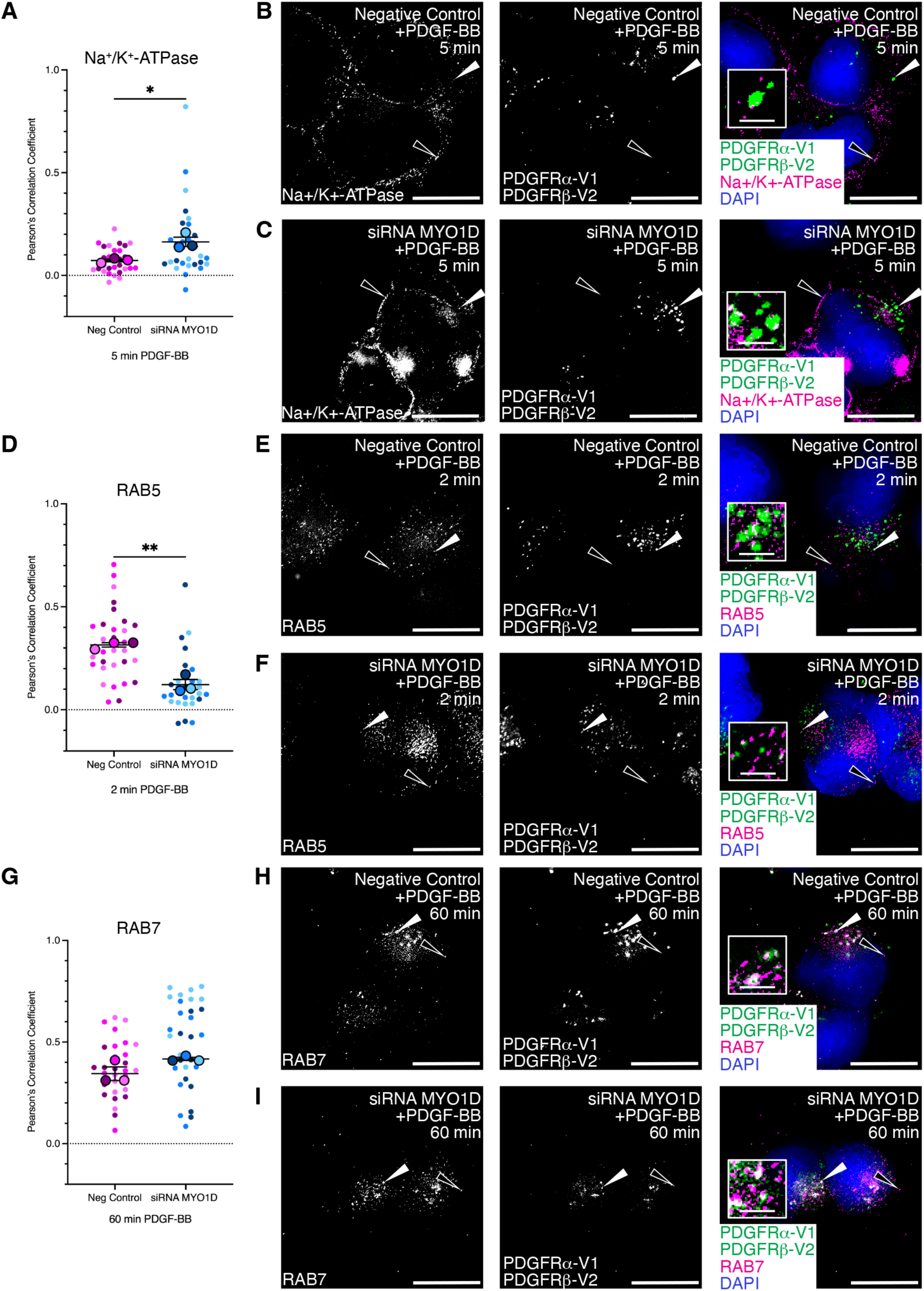
Knockdown of MYO1D leads to retention of PDGFRα/β heterodimers at the plasma membrane. (**A,D,G**) Scatter dot plots depicting Pearson’s correlation coefficient of Venus signal from the PDGFRα/β heterodimer cell line treated with Silencer Select negative control or Silencer Select siRNA against MYO1D with an anti-Na^+^/K^+^-ATPase antibody (A), an anti-RAB5 antibody (D) or an anti-RAB7 antibody (G) signal following PDGF-BB ligand stimulation for 5 min (A), 2 min (D) or 60 min (G). Data are mean±s.e.m. \**P*<0.05; \*\**P*<0.01 (two-tailed, unpaired *t*-test with Welch’s correction). Colored circles correspond to independent experiments. Summary statistics from biological replicates consisting of independent experiments (large circles) are superimposed on top of data from all cells; *n*=10 technical replicates across each of three biological replicates. (**B,C,E,F,H,I**) Na^+^/K^+^-ATPase antibody signal (white or magenta; B,C), RAB5 antibody signal (white or magenta; E,F) or APPL1 antibody signal (white or magenta; H,I) and/or Venus expression (white or green; B,C,E,F,H,I) as assessed by (immuno)fluorescence analysis of the PDGFRα/β heterodimer cell line. Insets in B, C, E, F, H and I are regions where white arrows are pointing. Nuclei were stained with DAPI (blue; B,C,E,F,H,I). White arrows denote colocalization; white outlined arrows denote lack of colocalization. Scale bars: 20 µm (main images), 3 µm (insets).

Further, knockdown of MYO1D lead to changes in intracellular signaling downstream of PDGFRα/β heterodimer activation. Specifically, phospho-ERK1/2 levels were increased upon treatment with siRNAs targeting *MYO1D* at all timepoints tested from 2 min to 4 h, with the largest difference between negative control (1.15±0.234 R.I.) and experimental siRNA treatment (1.51±0.257 R.I.) occurring at 5 min of ligand treatment (Fig. 7A to C). Alternatively, phospho-AKT levels were decreased upon treatment with siRNAs targeting *MYO1D* at the majority of timepoints tested, with the largest difference between negative control (1.90±0.276 R.I.) and experimental siRNA treatment (1.67±0.0927 R.I.) also occurring at 5 min of ligand treatment (Fig. 7D to F).

**Fig. 7.**
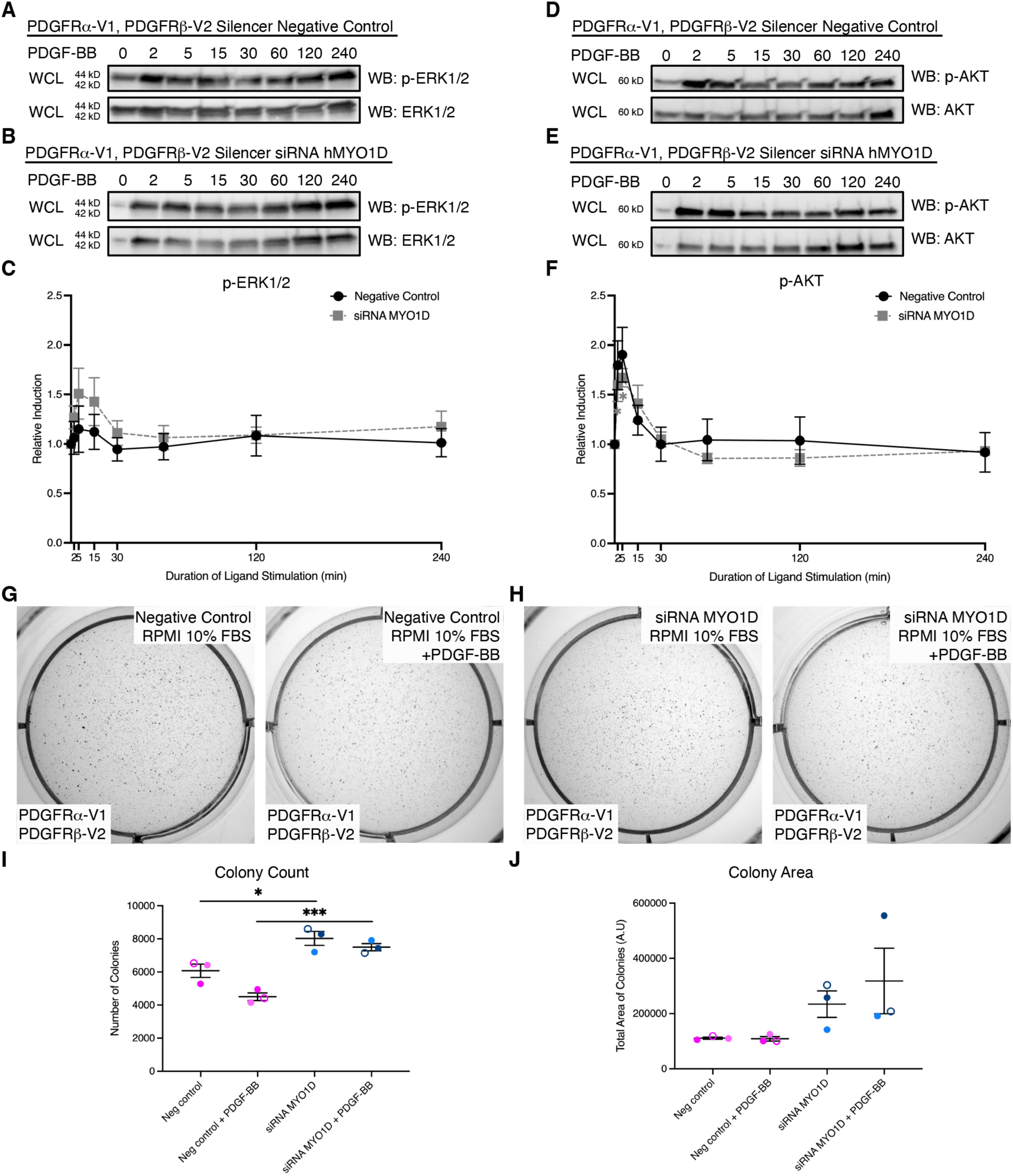
Knockdown of MYO1D leads to increased downstream phosphorylation of ERK1/2 and increased cell proliferation. (**A,B,D,E**) Western blot (WB) analysis of whole-cell lysates (WCL) from the PDGFRα/β heterodimer cell line treated with Silencer Select negative control (A,D) or Silencer Select siRNA against MYO1D (B,E) following a time course of PDGF-BB ligand stimulation from 2 min to 4 h with anti-phospho (p)-ERK1/2 (A,B) or anti-phospho (p)-AKT (D,E) antibodies. (**C,F**) Line graphs depicting quantification of band intensities from *n*=3 biological replicates as in A, B, D and E. Data are mean±s.e.m. \**P*<0.05 (two-tailed, ratio paired *t*-test). (**G,H**) Colony growth in soft agar anchorage-independent growth assays for the PDGFRα/β heterodimer cell line treated with Silencer Select negative control (G) or Silencer Select siRNA against MYO1D (H) after 10 days in RPMI growth medium in the absence or presence of PDGF-BB ligand in six-well plate wells. (**I,J**) Scatter dot plots depicting quantification of colony count (I) or colony area (J) from *n*=3 biological replicates as in G and H. Data are mean±s.e.m. \**P*<0.05; \*\*\**P*<0.001 (two-tailed, paired *t-*test within each siRNA treatment and a two-tailed, unpaired *t*-test with Welch’s correction between each siRNA treatment). Colored symbols correspond to independent experiments.

Finally, we assessed cell proliferation downstream of PDGFRα/β heterodimer activation upon knockdown of MYO1D. Soft agar anchorage-independent growth assays revealed that PDGFRαV1/βV2 cells treated with negative control (Fig. 7G) proliferated less than the same cells treated with siRNAs targeting *MYO1D* (Fig. 7H) in the absence (6.07×10^3^±4.00×10^2^ versus 8.03×10^3^±4.22×10^2^ colonies, respectively) and presence (4.50 ×10^3^±2.32×10^2^ versus 7.50×10^3^±2.21×10^2^ colonies, respectively) of PDGF-BB ligand (Fig. 7I and J), and significantly so as assessed by colony count (Fig. 7I). Taken together, these data demonstrate that knockdown of MYO1D leads to retention of PDGFRα/β heterodimers at the plasma membrane, further resulting in increased downstream phosphorylation of ERK1/2 and increased cell proliferation.

## DISCUSSION

In this study, we used the BiFC technique to visualize and purify PDGFRα/β heterodimers, demonstrating that this continues to be an invaluable approach for studying RTK dimer-specific properties (*24, 25, 44*). It is useful to compare our findings across the three PDGFR-BiFC cell lines, as the experimental parameters were identical. First, all three dimers exhibited peak dimerization following 5 min of ligand stimulation, with PDGFRα/β heterodimers exhibiting an intermediate speed of dimerization compared to the two homodimers (*24*). Further, the PDGFRα/β heterodimers had considerably reduced levels of autophosphorylation compared to either homodimer, particularly at early ligand stimulation timepoints (*24*). Next, PDGFRα/β heterodimers were internalized into early endosomes more quickly and dwelled at that location for longer periods of time than either homodimer (*24*). However, the extent of colocalization of the Venus signal from PDGFRα/β heterodimers and PDGFRβ homodimers with markers of late and recycling endosomes was comparable, indicating that these two dimers undergo degradation and recycling with similar dynamics and are less likely to be degraded and more likely to be recycled than PDGFRα homodimers (*24*). Perhaps most surprisingly, phospho-ERK1/2 levels were not sustained in response to PDGFRα/β heterodimer activation, and the phospho-AKT response, while reaching a higher relative induction than upon PDGFRα homodimer activation, was very transient in the PDGFRα/β heterodimer cell line upon PDGF-BB ligand treatment (*24*). Accordingly, PDGFRα/β heterodimer activation did not induce a proliferative response, unlike the robust proliferation observed upon PDGFRα and PDGFRβ homodimer activation (*24*).

One limitation of this study is that we cannot rule out that differences in expression levels of the PDGFRs across our three PDGFR-BiFC cell lines contribute to the disparities in results discussed above. Though we were able to identify a PDGFRα/β heterodimer clone with relatively equal expression of *PDGFRA* and *PDGFRB* among the hundreds of clones screened, the clone had reduced overall levels of these transcripts compared to our PDGFRα homodimer and PDGFRβ homodimer clones (*24*). In the future, it will be critical to compare the dynamics of the various PDGFRs at endogenous expression levels, ideally in a cell type in which there is evidence that all three dimers form, such as the craniofacial mesenchyme (*8*). In that setting, BiFC should continue to serve as a robust approach to both visualize and purify the individual PDGFR dimers. A second limitation of our work is that we cannot be sure that PDGFRα/β heterodimers that colocalize with Na^+^/K^+^-ATPase at 60 and 90 min of ligand treatment have been recycled. Live-cell imaging over such long timepoints is not possible in our system due to photobleaching of Venus. However, the combination of more photostable fluorescent proteins with the BiFC technique should prove a powerful approach to overcome this limitation in the future.

Our colocalization studies demonstrated that PDGFRα/β heterodimers are rapidly internalized into the signaling endosome subpopulation of early endosomes, where they dwell for extended lengths of time. The fact that PDGFRα/β heterodimers are not long-lived at the plasma membrane indicates that they are unlikely to generate a robust signaling response at this subcellular platform. Given that the signaling endosome marker APPL1 (*36*) has previously been shown to associate with the RTKs EGFR and TrkA and is required for Akt and Erk1/2 signaling downstream of these receptors (*45–48*), it was unexpected that PDGFRα/β heterodimers did not induce robust and sustained phospho-ERK1/2 and phospho-AKT signals from this site either. It is possible, however, that PDGFRα/β heterodimers may sequester APPL1, thereby preventing it from propagating signaling downstream of other RTKs. In fact, our results point to a similar role for PDGFRα/β heterodimers in serving as sinks that prevent PDGF-BB ligand from binding and activating PDGFRβ homodimers. While the biological significance of this potential function of PDGFRα/β heterodimers is unknown, it is tempting to speculate that heterodimer formation may serve to attenuate PDGFR signaling in instances of cell crowding and/or increased PDGF-BB ligand availability. Though assays comparing the binding affinity of PDGFRα/β heterodimers and PDGFRβ homodimers to PDGF-BB ligand are beyond the scope of this work, such studies would likely shed further light on the role of PDGFRα/β heterodimers.

Our mass spectrometry findings represent the first PDGFR dimer-specific interactome and will be a rich resource for the field. Of note, our screen revealed an enrichment for proteins that function in mRNA processing, and specifically mRNA splicing, in binding all three PDGFR dimers. PI3K has been identified as the main downstream effector of PDGFRα signaling during murine craniofacial development (*16, 49*). Interestingly, we had previously demonstrated in the context of the facial mesenchyme that phosphorylation targets of Akt downstream of PI3K-mediated PDGFRα homodimer signaling are similarly enriched for proteins involved in RNA processing (*49*) and further, that alternative RNA splicing is the predominant mechanism of gene expression regulation downstream of this signaling pathway (*50*). Taken together, these results indicate that proteins that function in mRNA processing engage with PDGFR signaling on multiple levels: through binding to the receptors and via phosphorylation by downstream effector(s). Moreover, our finding that PDGFRα homodimers interact with proteins that function in organization of the actomyosin cytoskeleton is in line with prior work demonstrating that PDGFRα homodimer signaling drives association of the transcription factor serum response factor (SRF) with the myocardin-related transcription factor-A (MRTFA) cofactor to upregulate expression of gene products that function within the actomyosin cytoskeleton (*51*). Finally, the fact that PDGFRα homodimers bound proteins that function in the regulation of signal transduction by p53 is consistent with our prior finding that PI3K-mediated PDGFRα homodimer signaling regulates p53-dependent intracellular pathways in the context of skeletal development (*49*). Though the full scope of PDGFRα/β heterodimer function remains to be determined, our mass spectrometry results suggest that activation of this dimer may ultimately contribute to nucleosome assembly and organization, and regulation of autophagy, immunoglobulin production and gene expression. Given our findings that PDGFR dimers are dynamically trafficked throughout the cell for several hours following ligand treatment (*24*), similar approaches coupling biochemical purification of the dimers followed by mass spectrometry at multiple timepoints following ligand stimulation should provide a more complete understanding of the PDGFR dimer-specific interactome in various subcellular compartments. Further, as varying concentrations of PDGF ligand have been shown to result in differences in the timing and extent of PDGFR internalization, phosphorylation of downstream effector molecules, proliferation and migration (*52*), analogous experiments to the ones performed here with varying PDGF ligand concentrations should determine whether differential engagement of interacting proteins contributes to these responses.

In this study, we identified MYO1D as a novel protein interacting with the PDGFRs and demonstrated that it preferentially binds to dimerized, phosphorylated PDGFRα/β heterodimers. We do not know at this time whether MYO1D binds PDGFRα and/or PDGFRβ in the heterodimer complex, nor at what site(s) in the receptor(s) this binding occurs. These questions will be the subject of future research. However, given that the heavy/light ratio in our mass spectrometry screen was also greater than 1 for the PDGFRα homodimers (1.73), we speculate that MYO1D binds the PDGFRα receptor within the heterodimer complex upon ligand treatment. We found that MYO1D contributes to the rapid internalization of PDGFRα/β heterodimers from the plasma membrane into early endosomes in HCC15 lung cancer cells. Interestingly, these results are in direct contrast to what has previously been demonstrated for EGFR, ERBB2 and ERBB4 in Caco2 colorectal cancer cells. For these members of the ERBB family, MYO1D was shown to bind the unphosphorylated receptors and anchor them in the plasma membrane (*43*). As such, siRNA-mediated knockdown of MYO1D resulted in decreased EGFR, ERBB2 and ERBB4 levels at the plasma membrane (*43*). It is possible that these discrepancies arise due to differing cellular contexts and/or expression levels of the various RTKs in their respective settings. Alternatively, these results may reflect opposing roles for MYO1D in regulating RTK localization and represent a means of divergence across RTK families.

Here, we demonstrated that PDGFRα/β heterodimers are rapidly internalized from the plasma membrane into early endosomes, resulting in an absence of phospho-ERK1/2 induction and a transient phospho-AKT response. Alternatively, retention of PDGFRα/β heterodimers at the plasma membrane via knockdown of MYO1D led to increased phospho-ERK1/2 levels and decreased phospho-AKT levels across multiple timepoints of ligand treatment, with the largest differences between negative control and experimental siRNA treatments observed at the relatively early timepoint of 5 min of ligand treatment. These findings indicate that ERK1/2 and AKT phosphorylation downstream of PDGFRα/β activation may primarily occur at the plasma membrane and the early endosome, respectively, immediately following ligand stimulation. Relatedly, our previous results demonstrated that inhibition of clathrin-mediated endocytosis of PDGFRβ homodimers via Dyngo-4a treatment resulted in increased phospho-ERK1/2 levels and decreased phospho-AKT levels at early timepoints of ligand stimulation (*24*), indicating that engagement of these signaling molecules occurs at the same subcellular platforms for PDGFRα/β heterodimers and PDGFRβ homodimers. Alternatively, we previously observed that Dyngo-4a treatment of the PDGFRα homodimer cell line had a negative effect on ERK1/2 and AKT phosphorylation, resulting in delayed and reduced peaks of activation (*24*). These results demonstrated that rapid internalization of PDGFRα homodimers plays an important role in the propagation of downstream signaling and suggested that this dimer primarily signals from early endosomes. Taken together, our findings support a model in which protein interactions and subcellular localization represent mechanisms by which specificity is introduced downstream of PDGFR activation to differentially propagate signaling and generate distinct cellular responses.

## MATERIALS AND METHODS

### Generation of the PDGFRα/β-BiFC-HCC15 and myr-Venus-HCC15 cell lines

The PDGFRαV1/αV2 homodimer and PDGFRβV1/βV2 homodimer cell lines were generated as previously described (*24*). For the PDGFRαV1/βV2 heterodimer cell line, PDGFRα-V1/pLVX-Puro (*24*) and PDGFRβ-V2/pLVX-Puro (*24*) lentiviral constructs (10 mg each) and packaging vectors pCMV-VSV-G (*53*) and pCMV-dR8.91 (*54*) (5 mg each) were transfected into HEK 293T/17 cells (American Type Culture Collection) using Lipofectamine LTX (Thermo Fisher Scientific, Waltham, MA, USA). For the myristoylated-Venus-HCC15 cell line, the pCAG:myr-Venus sequence (#32602, Addgene, Watertown, MA, USA) was amplified by PCR using the following primers (Integrated DNA Technologies, Inc., Coralville, IA, USA): 5’-GATAGGATCCGCCACCATG-3’ and 5’-CGTTTCTAGATTACTTGTACAGCTCG-3’. The sequence was cloned into the pLVX-Puro vector using BamHI and XbaI sites. The myr-Venus/pLVX-Puro lentiviral construct (10 mg) and packaging vectors pCMV-VSV-G and pCMV-dR8.91 (5 mg each) were transfected into HEK 293T/17 cells using Lipofectamine LTX. In each case, medium containing lentivirus was collected 48 h and 72 h following transfection and filtered using a 13 mm syringe filter with a 0.45 µm PVDF membrane (Thermo Fisher Scientific) following addition of 4 mg/ml polybrene (Sigma-Aldrich, St Louis, MO, USA). Lentiviral-containing medium was added to HCC15 lung cancer cells (see below) every 24 h for 2 days, and cells were subsequently grown in the presence of 2 µg/ml puromycin for 10 days. Individual Venus-positive cells were isolated on a Moflo XDP 100 cell sorter (Beckman Coulter Inc., Brea, CA, USA), following 5 min of stimulation with 10 ng/ml PDGF-BB (R&D Systems, Minneapolis, MN, USA) in the case of the PDGFRαV1/βV2 heterodimer cell line, and expanded to generate clonal cell lines. The final clone chosen for each cell line was confirmed by PCR amplification of the inserted sequence(s) from genomic DNA using primers found in Table S1. These PCR products were subcloned into the Zero Blunt TOPO vector (Thermo Fisher Scientific) and Sanger sequenced.

### PDGFR-BiFC-HCC15 and myr-Venus-HCC15 cell culture

The HCC15 lung cancer cell line was obtained from the laboratory of Dr. Lynn Heasley (University of Colorado Anschutz Medical Campus). The cell line was authenticated through short tandem repeat analysis and tested for mycoplasma contamination every 10 passages using the MycoAlert Mycoplasma Detection Kit (Lonza Group Ltd, Basel, Switzerland). PDGFR-BiFC and myr-Venus stable cells were cultured in RPMI growth medium [RPMI 1640 (Gibco, Thermo Fisher Scientific) supplemented with 100 U/ml penicillin (Gibco), 100 µg/ml streptomycin (Gibco) containing 10% FBS (Hyclone Laboratories Inc., Logan, UT, USA)] at 37°C in 5% CO_2_. Once the stable cell lines were established, they were split at a ratio of 1:5 for maintenance. PDGFRα/β heterodimer cells were used for experiments at passages 10-17, PDGFRα homodimer cells were used for experiments at passages 39-44, PDGFRβ homodimer cells were used for experiments at passages 48-53, and myr-Venus cells were used for experiments at passages 10-13. When serum starved, cells were grown in HITES medium [DMEM/F12 (Corning, Corning, NY, USA) supplemented with 0.1% bovine serum albumin (Fisher Scientific, Thermo Fisher Scientific), 10 mM β-estradiol (Sigma-Aldrich), 10 mM hydrocortisone (Sigma-Aldrich), 5 µg/ml insulin (Sigma-Aldrich), 100 U/ml penicillin (Gibco), 100 µg/ml streptomycin (Gibco), 1.2 mg/ml NaHCO_3_ (Santa Cruz Biotechnology, Inc., Dallas, TX, USA), 30 nM Na_3_SeO_3_ (Sigma-Aldrich) and 10 µg/ml apo-transferrin (Sigma-Aldrich)].

### qRT-PCR

Total RNA was isolated using the RNeasy Mini Kit (Qiagen, Germantown, MD, USA) according to the manufacturer’s instructions. First-strand cDNA was synthesized using a ratio of 2:1 random primers:oligo (dT) primer and SuperScript II RT (Invitrogen, Thermo Fisher Scientific) according to the manufacturer’s instructions. qRT-PCR was performed on a CFX Connect Real-Time PCR Detection System and analyzed with CFX Manager software (version 3.1; Bio-Rad Laboratories, Inc., Hercules, CA, USA). All reactions were performed with SYBR Select Master Mix (Applied Biosystems, Thermo Fisher Scientific), 300 nM primers (Integrated DNA Technologies, Inc.) and cDNA in a 20 µl reaction volume. PCR primers for qRT-PCR analyses can be found in Table S1. The following PCR protocol was used: step 1, 2 min at 50°C; step 2, 2 min at 95°C; step 3, 15 s at 95°C; step 4, 1 min at 60°C; repeat steps 3 and 4 for 39 cycles; step 5 (melting curve), 5 s per 0.5°C increment from 65°C to 95°C. All samples were run in triplicate and normalized against an endogenous internal control, *B2M*. qRT-PCR experiments were performed across three independent experiments, each using a separate passage of cells.

### Immunoprecipitations and western blotting

PDGFR-BiFC-HCC15 cells and myr-Venus-HCC15 cells were cultured as described above. To induce PDGFRα homodimer, PDGFRα/β heterodimer or PDGFRβ homodimer signaling, PDGFRα homodimer cells and PDGFRβ homodimer cells at ∼60–70% confluence or PDGFRα/β heterodimer cells at ∼70-80% confluence were serum starved for 24 h in HITES medium and stimulated with 10 ng/ml PDGF-AA (PDGFRα homodimer cells) or PDGF-BB (PDGFRα/β heterodimer and PDGFRβ homodimer cells) ligand (R&D Systems) for the indicated length of time. When applicable, cells were transfected with siRNA (see below) for 48 h before ligand stimulation. Protein lysates for immunoprecipitation were generated by resuspending cells in ice-cold GFP-Trap buffer [M-PER mammalian protein extraction reagent (Thermo Fisher Scientific), 1x complete Mini protease inhibitor cocktail (Roche, MilliporeSigma, Burlington, MA, USA), 1 mM PMSF, 10mM NaF, 1 mM Na_3_VO_4_, 25 mM β-glycerophosphate] and collecting cleared lysates by centrifugation at 13,400 *g* at 4°C for 20 min. For immunoprecipitations, cell lysates (500 µg for PDGFRα homodimers and PDGFRβ homodimers, 1 mg for PDGFRα/β heterodimers) were incubated with GFP-Trap agarose beads (Bulldog Bio, Inc., Portsmouth, NH, USA) for 1 h at 4°C. Beads were washed three times with ice-cold GFP-Trap buffer and the precipitated proteins were eluted with Laemmli buffer containing 10% β-mercaptoethanol, heated for 10 min at 100°C and separated by SDS-PAGE. When applicable, supernatant from immunoprecipitation experiments was incubated with primary antibody overnight at 4°C followed by incubation with 20 µL of Protein A/G Plus-agarose beads (Santa Cruz Biotechnology, Inc.) for 2 h at 4°C the following day. Beads were washed five times with ice-cold GFP-Trap buffer and the precipitated proteins were eluted with Laemmli buffer containing 10% β-mercaptoethanol, heated for 5 min at 95°C, and separated by SDS-PAGE. For western blotting analysis of whole-cell lysates, protein lysates were generated by resuspending cells in ice-cold GFP-Trap buffer and collecting cleared lysates by centrifugation at 13,400 *g* at 4°C for 20 min. Laemmli buffer containing 10% β-mercaptoethanol was added to the lysates, which were heated for 5 min at 100°C. Proteins were subsequently separated by SDS-PAGE. Western blot analysis was performed according to standard protocols using horseradish peroxidase-conjugated secondary antibodies. Blots were imaged using a ChemiDoc XRS+ (Bio-Rad Laboratories, Inc.) or a ChemiDoc (Bio-Rad Laboratories, Inc.). The following antibodies were used for western blotting: PDGFRα (1:1000; D13C6; 5241; Cell Signaling Technology, Inc., Danvers, MA, USA); PDGFRβ (1:1000; 28E1; 3169; Cell Signaling Technology, Inc.); β-tubulin (1:1000; E7; E7; Developmental Studies Hybridoma Bank, Iowa City, IA, USA); Lamin B1 (1:1000; D4Q4Z, 12586; Cell Signaling Technology Inc.); phospho-PDGFRα (Tyr 849)/PDGFRβ (Tyr857) (1:1000; C43E9; 3170; Cell Signaling Technology, Inc.); phospho-ERK1/2 (Thr202/Tyr204) (1:1000; 9101; Cell Signaling Technology, Inc.); ERK1/2 (1:1000; 9102; Cell Signaling Technology, Inc.); phospho-AKT (Ser473) (1:1000; 9271; Cell Signaling Technology Inc.); AKT (1:1000; 9272; Cell Signaling Technology Inc.); Myosin ID (1:500; H-1; sc-515292; Santa Cruz Biotechnology, Inc); GAPDH (1:50000; 1E6D9; 60004-1-Ig; Proteintech Group Inc, Rosemont, IL, USA); horseradish peroxidase-conjugated goat anti-rabbit IgG (1:20,000; 111035003; Jackson ImmunoResearch Inc., West Grove, PA, USA); and horseradish peroxidase-conjugated goat anti-mouse IgG (1:20,000; 115035003; Jackson ImmunoResearch Inc.). Quantifications of signal intensity were performed with ImageJ software (version 1.53t, National Institutes of Health, Bethesda, MD, USA). Relative dimerized PDGFR levels were determined by normalizing GFP-Trap immunoprecipitated PDGFR levels to total PDGFR levels. Relative phospho-PDGFR levels were determined by normalizing to total PDGFR levels, except for ligand sink experiments in which relative phospho-PDGFR levels were determined by normalizing to unstimulated phospho-PDGFR signals. Relative phospho-ERK1/2 levels were determined by normalizing to total ERK1/2 levels. Relative phospho-AKT levels were determined by normalizing to total AKT levels. When applicable, statistical analyses were performed with Prism 9 (GraphPad Software Inc., San Diego, CA, USA) using a two-tailed, ratio paired *t*-test within each cell line (comparing individual ligand treatment timepoint values to the no ligand 0 min timepoint value) or a two-tailed, unpaired *t*-test with Welch’s correction between each treatment. Immunoprecipitation and western blotting experiments were performed across three independent experiments.

### Fluorescence analysis

For fluorescence intensity and marker colocalization experiments, cells were seeded onto glass coverslips coated with 5 µg/ml human plasma fibronectin purified protein (MilliporeSigma) at a density of 80,000 cells, 100,000 cells and 40,000 cells per 24-well plate well for the PDGFRα homodimer cell line, the PDGFRα/β heterodimer cell line and the PDGFRβ homodimer cell line, respectively, in RPMI growth medium. After 24 h, cells were washed with 1x phosphate buffered saline (PBS) and serum starved in HITES medium. HITES medium was replaced 23 h later. After 54 min, coverslips were photobleached for 1 min with an Axio Observer 7 fluorescence microscope (Carl Zeiss Microscopy LLC, White Plains, NY, USA) using the 2.5x objective and 488 nm laser. Cells were allowed to recover for 5 min and were treated with 10 ng/ml PDGF-AA (PDGFRα homodimer cells) or PDGF-BB (PDGFRα/β heterodimer and PDGFRβ homodimer cells) ligand (R&D Systems) for the indicated amount of time. When applicable, cells were transfected with siRNA (see below) for 48 h before ligand stimulation. Cells were fixed in 4% paraformaldehyde (PFA) in PBS with 0.1% Triton X-100 for 10 min and washed in PBS. Cells were blocked for 1 h in 5% normal donkey serum (Jackson ImmunoResearch Inc.) in PBS and incubated overnight at 4°C in primary antibody diluted in 1% normal donkey serum in PBS. After washing in PBS, cells were incubated in Alexa Fluor 546-conjugated donkey anti-rabbit secondary antibody (1:1000; A21206; Invitrogen) or Alexa Fluor 546-conjugated donkey anti-mouse secondary antibody (1:1000; A10036; Invitrogen) diluted in 1% normal donkey serum in PBS with 2 µg/ml DAPI (Sigma-Aldrich) for 1 h. Cells were mounted in VECTASHIELD HardSet Antifade Mounting Medium (Vector Laboratories, Inc., Burlingame, CA, USA) and photographed using an Axiocam 506 mono digital camera (Carl Zeiss Microscopy LLC) fitted onto an Axio Observer 7 fluorescence microscope (Carl Zeiss Microscopy LLC) with the 63x oil objective with a numerical aperture of 1.4 at room temperature. The following antibodies were used for immunofluorescence analysis: PDGFRα (1:1000; D13C6; 5241; Cell Signaling Technology, Inc.); PDGFRβ (1:1000; 28E1; 3169; Cell Signaling Technology, Inc.); Na^+^/K^+^-ATPase (1:500; EP1845Y; ab76020; Abcam); RAB5 (1:200; C8B1; 3547; Cell Signaling Technology Inc.); APPL1 (1:200; D83H4; 3858; Cell Signaling Technology Inc.); EEA1 (1:200; C45B10; 3288; Cell Signaling Technology Inc.); RAB7 (1:100; D95F2; 9367; Cell Signaling Technology Inc.); RAB4 (1:200; ab13252; Abcam); RAB11 (1:100; D4F5; 5589; Cell Signaling Technology Inc.) and Myosin ID (1:500; H-1; sc-515292; Santa Cruz Biotechnology, Inc). For fluorescence intensity measurements, three independent trials, or biological replicates, were performed. For each biological replicate, at least 38 technical replicates consisting of individual cells were imaged with *Z*-stacks (0.24 µm between *Z*-stacks with a range of 2-8 *Z*-stacks). For marker colocalization experiments, three independent trials, or biological replicates, were performed. For each biological replicate, at least 10 technical replicates consisting of individual cells were imaged with *Z*-stacks (0.24 µm between *Z*-stacks with a range of 2-8 *Z*-stacks) per timepoint. Images were deconvoluted using ZEN Blue software (Carl Zeiss Microscopy LLC) using the ‘Better, fast (Regularized Inverse Filter)’ setting. For all images, extended depth of focus was applied to *Z*-stacks using ZEN Blue software (Carl Zeiss Microscopy LLC) to generate images with the maximum depth of field. For fluorescence intensity measurements, background was subtracted using rolling background subtraction with a radius of 30 pixels using Fiji software (version 2.1.0/1.53c). A region of interest (ROI) was drawn around each Venus-positive cell, and integrated density was measured and recorded as the fluorescence intensity. For marker colocalization measurements, an ROI was drawn around each Venus-positive cell in the corresponding Cy3 (marker) channel using Fiji software (version 2.1.0/1.53c). For each image with a given ROI, the Cy3 channel and the EGFP channel were converted to 8-bit images. Colocalization was measured using the Colocalization Threshold function, where the rcoloc value [Pearson’s correlation coefficient (PCC)] was used in statistical analysis. When applicable, statistical analyses were performed on the average values from each biological replicate with Prism 9 (GraphPad Software Inc.) using a two-tailed, paired *t*-test within each cell line (comparing individual ligand treatment timepoint values to the no ligand 0 min timepoint value) or a two-tailed, unpaired *t*-test with Welch’s correction between each cell line, siRNA treatment or non-0 min ligand treatment timepoints.

### Anchorage-independent growth assays

For measurement of anchorage-independent cell growth, 25,000 cells were suspended in 1.5 ml RPMI growth medium with 0.35% Difco Agar Noble (Becton, Dickinson and Company, Franklin Lakes, NJ) and overlaid on a base layer containing 1.5 ml RPMI growth medium with 0.5% Difco Agar Noble (Becton, Dickinson and Company) in six-well plate wells. A feeding layer of 2 ml RPMI growth medium supplemented with 10 ng/ml PDGF-BB ligand (R&D Systems) and/or 0.01 uM siRNA (see below) was added on top of the agar and replaced every day. Plates were incubated at 37°C in 5% CO_2_ for 10 days, and viable colonies were stained overnight with 1 mg/ml Nitrotetrazoleum Blue (Sigma-Aldrich) in PBS at 37°C. Following a second overnight incubation at 4°C, wells from each independent trial were photographed using a COOLPIX S600 digital camera (Nikon Inc., Melville, NY, USA). Images were made binary using Adobe Photoshop (version 25.0.1; Adobe Inc., San Jose, CA, USA), and colony number and area were quantified using MetaMorph imaging software (Molecular Devices, LLC, San Jose, CA, USA). Statistical analyses were performed on values from each of three independent trials, or biological replicates, with Prism 9 (GraphPad Software Inc.) using a two-tailed, paired *t*-test within each cell line or siRNA treatment and a two-tailed, unpaired *t*-test with Welch’s correction between each cell line or siRNA treatment.

### Mass spectrometry

The following amino acids were used for stable isotope labeling: unlabeled (light) L-LYSINE:2HCL, unlabeled (light) L-ARGININE:HCL, stable isotope-labeled (heavy) L-LYSINE:2HCL (13C6, 99%;15N2, 99%) and stable isotope-labeled (heavy) L-ARGININE:HCL (13C6, 99%; 15N4, 99%) (Cambridge Isotope Labs, Inc. Andover, MA). Light SILAC medium was generated by adding unlabeled L-LYSINE:2HCL (final concentration 146.0 mg/mL) and unlabeled L-ARGININE:HCL (final concentration 42.0 mg/mL) to RPMI SILAC growth medium [RPMI 1640 SILAC (Gibco, Thermo Fisher Scientific) supplemented with 100 U/ml penicillin (Gibco), 100 µg/ml streptomycin (Gibco) containing 10% FBS (Hyclone Laboratories Inc.)]. Heavy isotope SILAC medium was generated by adding heavy labeled L-LYSINE:2HCL (13C6, 99%;15N2, 99%) (final concentration 151.3 mg/mL) and heavy labeled L-ARGININE:HCL (13C6, 99%; 15N4, 99%) (final concentration 44.0 mg/mL) in RPMI SILAC growth medium. Each SILAC medium was filtered through a 0.22-um syringe filter. Each PDGFR-BiFC cell line was cultured in duplicate in light SILAC medium and separately in heavy SILAC medium for at least 6 cell passages to achieve >95% labeling. PDGFRα homodimer cells at ∼75% confluence, PDGFRα/β heterodimer cells at ∼70-80% confluence and PDGFRβ homodimer cells at ∼65% confluence were serum starved for 24 h in HITES medium. Cells grown in light SILAC medium were left unstimulated and cells grown in heavy SIILAC medium were stimulated with 10 ng/ml PDGF-AA (PDGFRα homodimer cells) or 10 ng/ml of PDGF-BB (PDGFRα/β heterodimer and PDGFRβ homodimer cells) ligand for 5 minutes. Myr-Venus cells were cultured in light SILAC medium, serum starved at ∼70% confluence for 24 h in HITES medium and left unstimulated. Protein lysates for immunoprecipitation were generated by resuspending cells in ice-cold GFP-Trap buffer and collecting cleared lysates by centrifugation at 13,400 *g* at 4°C for 20 min. Before immunoprecipitation, light SILAC samples were combined with heavy SILAC samples in a 1:1 ratio per cell line. For immunoprecipitations, cell lysates (500 µg for myr-Venus, PDGFRα homodimers and PDGFRβ homodimers, 1 mg for PDGFRα/β heterodimers) were incubated with GFP-Trap agarose beads (Bulldog Bio, Inc.) for 1 h at 4°C. Beads were washed three times with ice-cold GFP-Trap buffer.

To prepare for trypsin digestion, beads were resuspended with 8 M urea, 50 mM ammonium bicarbonate. Tris (2-carboxyethyl)phosphine (TCEP) was added to a final concentration of 10 mM and samples were incubated at 30°C for 1 h. Alkylation was performed by adding chloroacetamide (CAA) to 25 mM and incubating in the dark at room temperature for 30 min. Following alkylation, urea was diluted to 1 M and proteins digested overnight with modified sequencing grade trypsin (Promega, Madison, WI, USA) at a 1:50 enzyme:protein ratio. Beads were then centrifuged for 15 min at 18,000 *g* and supernatant digest solution was collected. Digested peptides were then desalted using Pierce C18 spin tips (Thermo Fisher Scientific) according to the manufacturer’s instructions.

Digested peptides were loaded onto individual Evotips (Evosep Biosystems, Odense, Denmark) following the manufacturer’s instructions and separated on an Evosep One chromatography system (Evosep Biosystems) using a Pepsep column (150 μm inter diameter, 15 cm) packed with ReproSil C18 1.9 μm, 120Å resin. Samples were analyzed using the instrument default “30 samples per day” LC gradient. The system was coupled to the timsTOF Pro mass spectrometer (Bruker Daltonics, Bremen, Germany) via the nano-electrospray ion source (Captive Spray, Bruker Daltonics). The mass spectrometer was operated in PASEF mode. The ramp time was set to 100 ms and 10 PASEF MS/MS scans per topN acquisition cycle were acquired. MS and MS/MS spectra were recorded from m/z 100 to 1700. The ion mobility was scanned from 0.7 to 1.50 Vs/cm2. Precursors for data-dependent acquisition were isolated within ± 1 Th and fragmented with an ion mobility-dependent collision energy, which was linearly increased from 20 to 59 eV in positive mode. Low-abundance precursor ions with an intensity above a threshold of 500 counts but below a target value of 20000 counts were repeatedly scheduled and otherwise dynamically excluded for 0.4 min.

Data was searched using PEAKS X Pro version 10.6 (Bioinformatics Solutions, Inc., Ontario, Canada). Precursor tolerance was set to ±25 ppm and fragment tolerance was set to ±0.2 Da. Data was searched against SwissProt (20,379 sequences) restricted to *Homo sapiens*. Each fraction was searched independently using a fully specific trypsin cleavage definition allowing for 2 missed cleavages and a maximum of 5 modifications per peptide. Fixed modifications were set as carbamidomethyl (C). Variable modifications were set as oxidation (M), acetylation (K, protein N-term), and phosphorylation (STY). SILAC isotope modifications 13C(6) 15N(4) (R) and 13C(6) 15N(2) (K) were also set as variable modifications. Results were filtered to 1% FDR at the peptide and protein level. SILAC quantification was performed using a “Quantification” task in PEAKS X Pro set for SILAC 2-plex (R10, K8) analysis. Quantification retention time range was set to 1.0 minute, with an ion mobility range of 0.05 1/k0. Heavy conditions were set to include 100% R(+10.01) and K(+8.01) modifications. All results were filtered to 1% FDR. Auto-normalization was applied to the quantification results within PEAKS before further processing.

### siRNA-mediated knockdown of Myosin ID

PDGFRα/β heterodimer cells at ∼60% confluence were transfected with Silencer Select Negative Control No. 1 siRNA (4390843; Invitrogen, Thermo Fisher Scientific) and separately two Silencer Select siRNAs targeting human Myosin ID (4392420; ID s9203 and s9204; Invitrogen, Thermo Fisher Scientific) at a final concentration of 0.01 uM using Lipofectamine RNAiMAX (Invitrogen, Thermo Fisher Scientific) according to the manufacturer’s instructions for the indicated amount of time.

## Supplementary Materials

**Fig. S1. Validation of a PDGFRα/β-BiFC stable cell line.**

**Fig. S2. PDGFRα/β heterodimers are ultimately trafficked for degradation and recycling.**

**Fig. S3. PDGFRα/β heterodimers may serve as ligand sinks.**

**Fig. S4. Gene ontology analysis of proteins that interact with the various PDGFR dimers.**

**Fig. S5. *MYO1D* expression is upregulated in the PDGFRα/β heterodimer cell line.**

**Fig. S6. Knockdown of MYO1D in the PDGFRα/β heterodimer cell line.**

**Table S1. PCR primers for confirmation of sequence integration and qRT-PCR analyses.**

## Acknowledgments

We thank Damian Garno, Jessica Johnston, Daniel Fuhr and Erin Binne for technical assistance. Cell sorting was performed at the University of Colorado Cancer Center Flow Cytometry Shared Resource with assistance from Dr. Dmitry Baturin. Mass spectrometry was performed at the University of Colorado School of Medicine Proteomics Core Facility with assistance from Sean Maroney. We are grateful to the laboratory of Dr. Rytis Prekeris for pCAG:myr-Venus, lentiviral and packaging plasmids and the laboratory of Dr. Lynn Heasley for HCC15 cells and advice on anchorage-independent growth assays. We thank members of the Fantauzzo laboratory for their critical comments on the manuscript.

## Author Contributions

Conceptualization: MBC, MAR, KAF; Methodology: MBC, MAR, KAF; Formal analysis: MBC, MCM, KAF; Investigation: MBC, MAR, MCM, AN, KAF; Writing – Original Draft: MBC, KAF; Writing – Review & Editing: MAR, MCM, AN, EDL; Visualization: MBC, EDL, KAF; Supervision: KAF; Funding acquisition: MBC, KAF.

