## Supplementary Materials for "PDGFRα/β heterodimer activation negatively affects downstream ERK1/2 signaling and cellular proliferation"

Maria B. Campaña *et al.*

**The PDF file includes:**

Figs. S1 to S6
Table S1

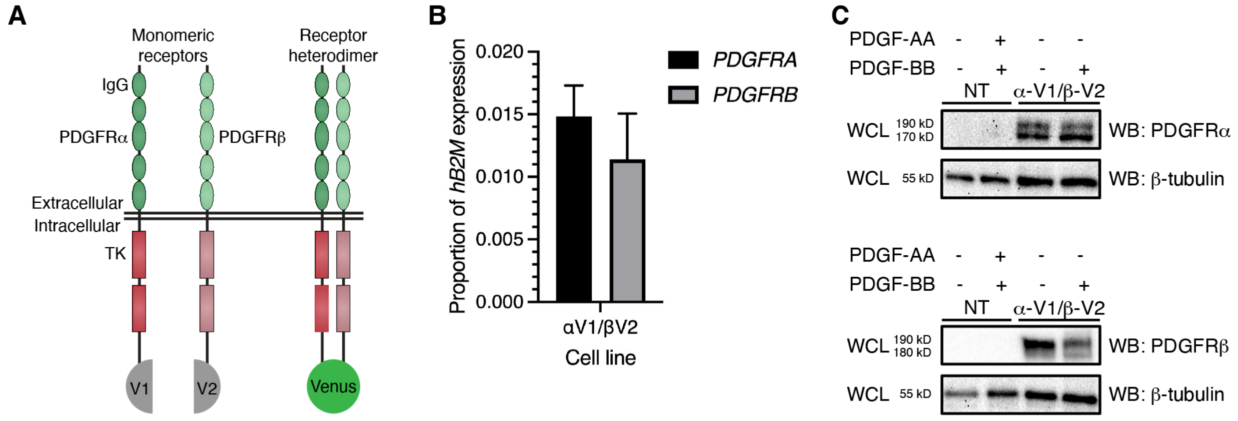


**Fig. S1. Validation of a PDGFRα/β-BiFC stable cell line.** (**A**) Schematic of PDGFRα and PDGFRβ with five extracellular immunoglobulin (IgG) domains and intracellular split tyrosine kinase (TK) domains fused to the non-fluorescent N-terminal (V1) and C-terminal (V2) fragments of Venus, respectively. Upon receptor dimerization, a functional Venus protein is generated. (**B**) Bar graph depicting *PDGFRA* and *PDGFRB* expression in the PDGFRα/β heterodimer cell line as assessed by quantitative RT-PCR. Data are mean±s.e.m. *n*=3 biological replicates. (**C**) Western blot (WB) analysis of whole-cell lysates (WCL) from non-transduced (NT) HCC15 cells (left) and PDGFRα/β heterodimer cells (right) in the absence or presence of PDGF ligand for 15 min with anti-PDGFRα (top) and anti-PDGFRβ (bottom) antibodies. *n*=2 biological replicates.

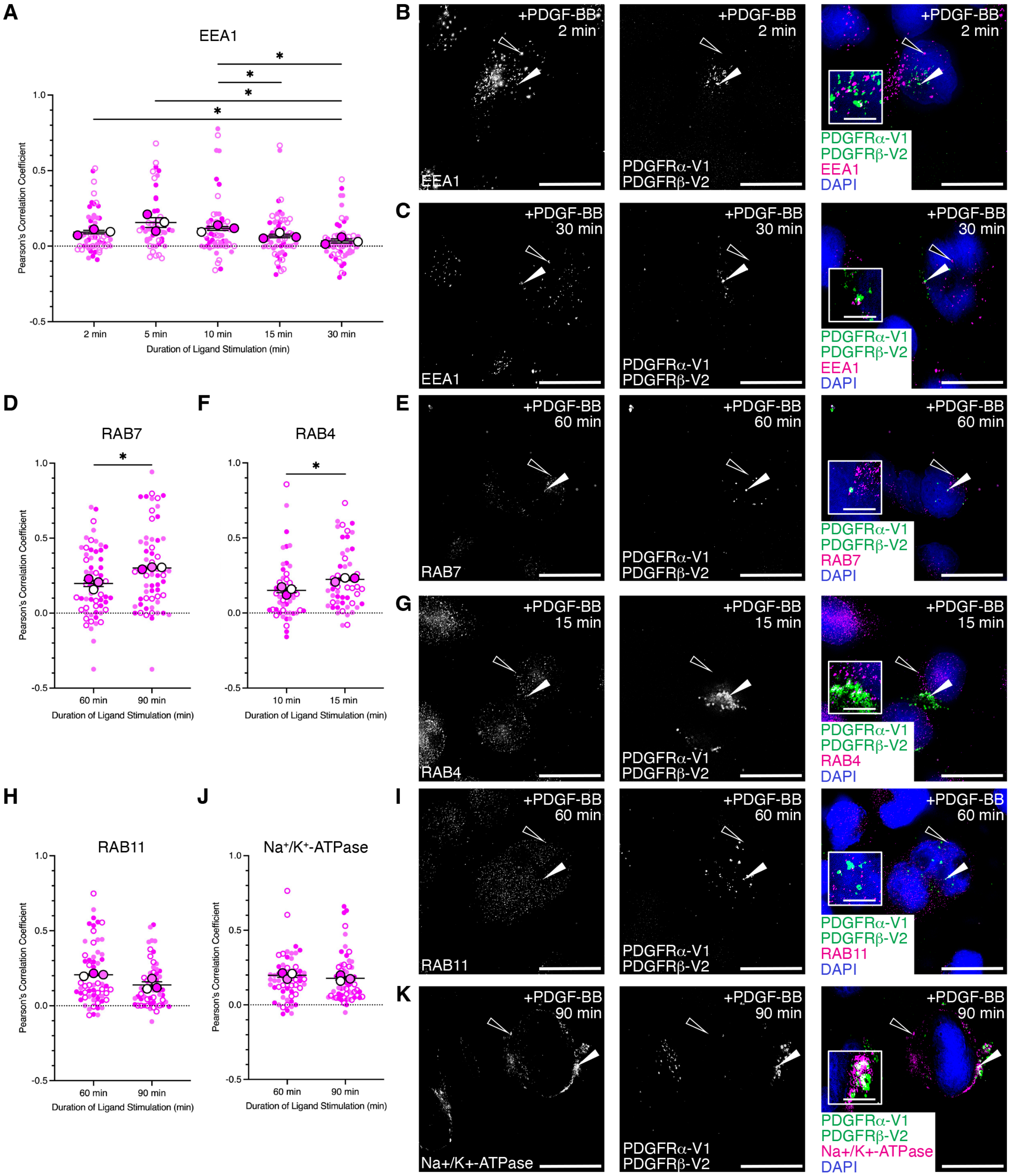


**Fig. S2. PDGFRα/β heterodimers are ultimately trafficked for degradation and recycling.** (**A,D,F,H,J**) Scatter dot plots depicting Pearson's correlation coefficient of the PDGFRα/β heterodimer cell line Venus signal with an anti-EEA1 antibody (A), an anti-RAB7 antibody (D), an anti-RAB4 antibody (F), an anti-RAB11 antibody (H) or an anti-$Na^+/K^+$-ATPase antibody (J) signal following PDGF-BB ligand stimulation from 2–30 min (A), 60-90 min (D,H,J) or 10-15 min (F). Data are mean±s.e.m. *$P$<0.05 (two-tailed, unpaired $t$-test with Welch's correction).

Colored circles correspond to independent experiments. Summary statistics from biological replicates consisting of independent experiments (large circles) are superimposed on top of data from all cells; $n \geq 15$ technical replicates across each of three biological replicates. (**B,C,E,G,I,K**) EEA1 antibody signal (white or magenta; B,C), RAB7 antibody signal (white or magenta; E), RAB4 antibody signal (white or magenta; G), RAB11 antibody signal (white or magenta; I) or $Na^+/K^+$-ATPase antibody signal (white or magenta; K) and/or Venus expression (white or green; B,C,E,G,I,K) as assessed by (immuno)fluorescence analysis of the PDGFRα/β heterodimer cell line. Insets in B, C, E, G, I and K are regions where white arrows are pointing. Nuclei were stained with DAPI (blue; B,C,E,G,I,K). White arrows denote colocalization; white outlined arrows denote lack of colocalization. Scale bars: 20 µm (main images), 3 µm (insets).

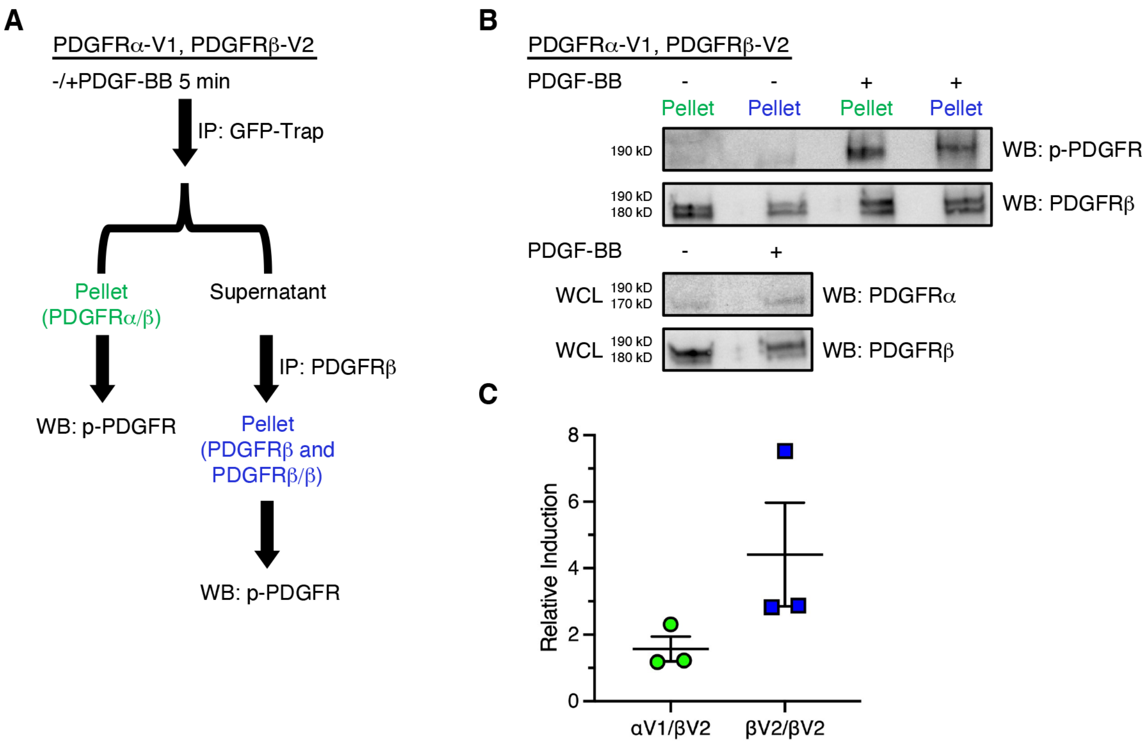


**Fig. S3. PDGFRα/β heterodimers may serve as ligand sinks.** (**A**) Schematic depicting experimental workflow to test whether PDGF-BB ligand could active PDGFRβ homodimers in the PDGFRα/β heterodimer cell line. (**B**) Immunoprecipitation (IP) of PDGFRα/β heterodimers with GFP-Trap nanobody (green) or immunoprecipitation of PDGFRβ monomers and PDGFRβ homodimers from the above supernatant with an anti-PDGFRβ antibody (blue) from cells that were unstimulated or treated with PDGF-BB ligand for 5 min followed by western blotting (WB) with an anti-phospho (p)-PDGFR antibody. WCL, whole-cell lysates. (**C**) Scatter dot plot depicting quantification of band intensities from *n*=3 biological replicates as in B. Data are mean±s.e.m.

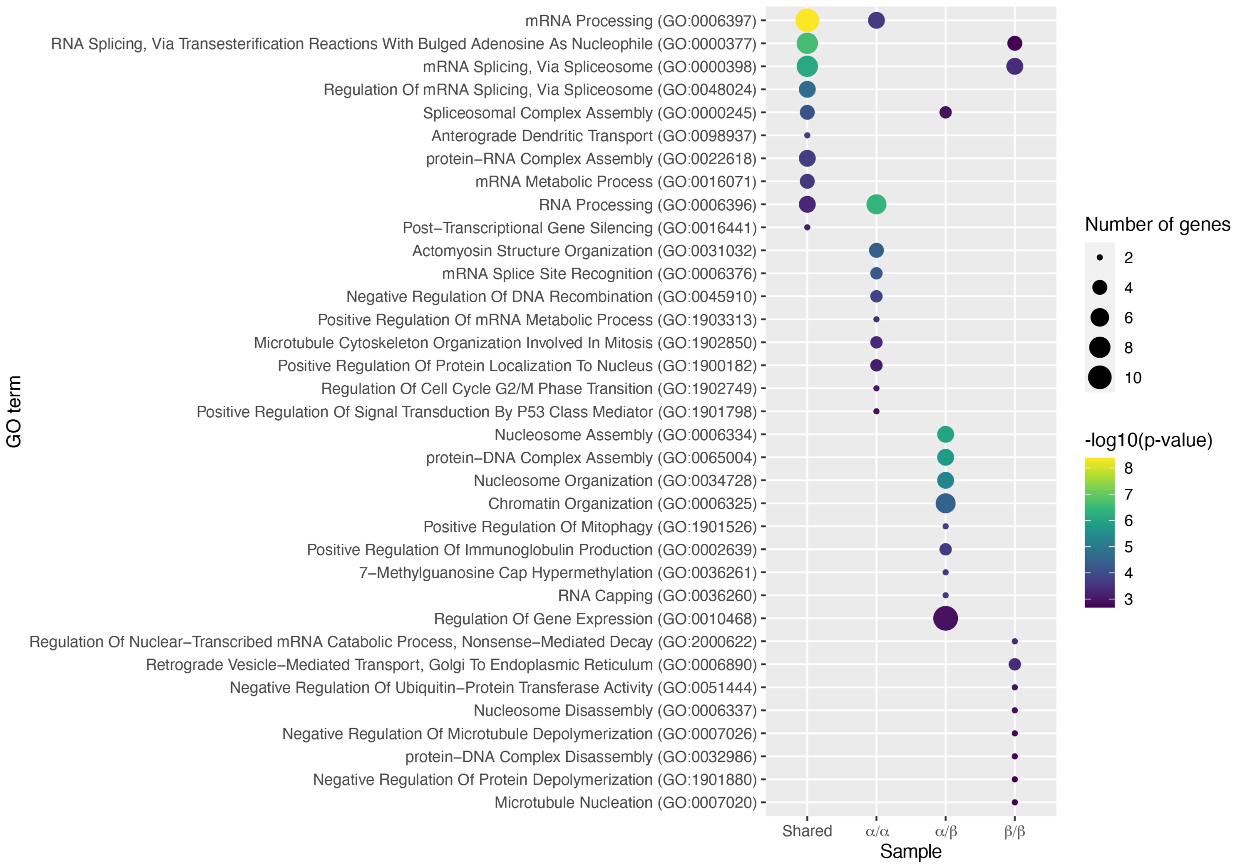


**Fig. S4. Gene ontology analysis of proteins that interact with the various PDGFR dimers.** Bubble plot depicting top ten biological process gene ontology (GO) terms for genes encoding the proteins that commonly interact with all three PDGFR dimers, or uniquely interact with PDGFRα homodimers, PDGFRα/β heterodimers or PDGFRβ homodimers as detected by mass spectrometry. Sizes correspond to number of genes; colors correspond to -log10(p-value).

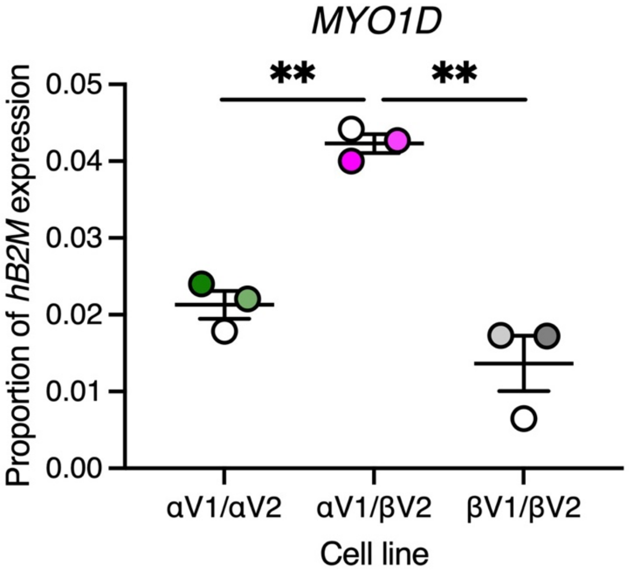


**Fig. S5. *MYO1D* expression is upregulated in the PDGFRα/β heterodimer cell line.** Scatter dot plot depicting *MYO1D* expression in the PDGFRα homodimer, PDGFRα/β heterodimer and PDGFRβ homodimer cell lines as assessed by quantitative RT-PCR. Data are mean±s.e.m. $^{**}P<0.01$ (two-tailed, unpaired *t*-test with Welch's correction). Colored circles correspond to independent experiments. Lighter circles correspond to later passages of cells. $n=3$ biological replicates.

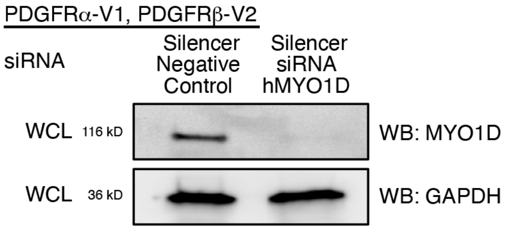


**Fig. S6. Knockdown of MYO1D in the PDGFRα/β heterodimer cell line.** Western blot (WB) analysis of whole-cell lysates (WCL) from cells transfected with a Silencer Select negative control siRNA or two Silencer Select siRNAs targeting *MYO1D* for 48 h. *n*=1 biological replicate.

**Table S1. PCR primers for confirmation of sequence integration and qRT-PCR analyses.**

| | |
|---|---|
| Common Forward Primer 1 | 5’-GGAGTTCCGCGTTACATAAC-3’ |
| PDGFRα Reverse 1 | 5’-GTTGGCCAAAATAGTCCAGG-3’ |
| PDGFRβ Reverse 1 | 5’-CTGAGATCACCACCACCTTA-3’ |
| PDGFRα Forward 2 | 5’-GCCGCTTCCTGATATTGAGT-3’ |
| PDGFRα Reverse 2 | 5’-GAATTATCTAGAGTCGCGGG-3’ |
| PDGFRβ Forward 2 | 5’-CTGCAGAGACCTCAAAAGGT-3’ |
| PDGFRβ Reverse 2 | 5’-CCAGACTGCCTTGGGAAAAG-3’ |
| Myr-Venus Reverse | 5’-CTAGATTACTTGTACAGCTCGTC-3’ |
| *B2M* Forward | 5’-CTACTCTCTCTTTCTGGCCT-3’ |
| *B2M* Reverse | 5’-GACAAGTCTGAATGCTCCAC-3’ |
| *PDGFRA* Forward | 5’-GAAGAGACCCTCCTTTTACC-3’ |
| *PDGFRA* Reverse | 5’-CTTCAGCTTGTCTTCCTCGT-3’ |
| *PDGFRB* Forward | 5’- CAATGCCATCAAACGGGGTT-3’ |
| *PDGFRB* Reverse | 5’- CACTCCTCAGAAACTCCTCA-3’ |
| *MYO1D* Forward | 5’-GTGACACGCCTCTTATTGAG-3’ |
| *MYO1D* Reverse | 5’-GATGCGAGTAACGATCCAAC-3’ |
